# Fonio millet genome unlocks African orphan crop diversity for agriculture in a changing climate

**DOI:** 10.1101/2020.04.11.037671

**Authors:** Michael Abrouk, Hanin Ibrahim Ahmed, Philippe Cubry, Denisa Šimoníková, Stéphane Cauet, Jan Bettgenhaeuser, Liubov Gapa, Yveline Pailles, Nora Scarcelli, Marie Couderc, Leila Zekraoui, Nagarajan Kathiresan, Jana Čížková, Eva Hřibová, Jaroslav Doležel, Sandrine Arribat, Hélène Bergès, Jan J. Wieringa, Mathieu Gueye, Ndjido A. Kane, Christian Leclerc, Sandrine Causse, Sylvie Vancoppenolle, Claire Billot, Thomas Wicker, Yves Vigouroux, Adeline Barnaud, Simon G. Krattinger

**Affiliations:** Center for Desert Agriculture, Biological and Environmental Science & Engineering Division (BESE), King Abdullah University of Science and Technology (KAUST), Thuwal, Saudi Arabia; DIADE, Univ Montpellier, IRD, Montpellier, France; Institute of Experimental Botany of the Czech Academy of Sciences, Centre of the Region Hana for Biotechnological and Agricultural Research, Olomouc, Czech Republic; CNRGV Plant Genomics Center, INRAE, Toulouse, France; Supercomputing Core Lab, King Abdullah University of Science and Technology (KAUST), Thuwal, Saudi Arabia; Inari Agriculture, One Kendall Square Building 600/700 Cambridge, MA 02139; Naturalis Biodiversity Center, Leiden, the Netherlands; Laboratoire de Botanique, Département de Botanique et Géologie, IFAN Ch. A. Diop/UCAD, Dakar, Senegal; Senegalese Agricultural Research Institute, Dakar, Senegal; Laboratoire Mixte International LAPSE, Dakar, Senegal; CIRAD, UMR AGAP, Montpellier, France; AGAP, Université de Montpellier, Cirad, INRAE, Institut Agro, Montpellier, France; Department of Plant and Microbial Biology, University of Zurich, Switzerland

**Author notes:** These authors contributed equally: Michael Abrouk, Hanin Ibrahim Ahmed, Philippe Cubry.

## Abstract

Sustainable food production in the context of climate change necessitates diversification of agriculture and a more efficient utilization of plant genetic resources. Fonio millet (*Digitaria exilis*) is an orphan African cereal crop with a great potential for dryland agriculture. Here, we established high-quality genomic resources to facilitate fonio improvement through molecular breeding. These include a chromosome-scale reference assembly and deep re-sequencing of 183 cultivated and wild *Digitaria* accessions, enabling insights into genetic diversity, population structure, and domestication. Fonio diversity is shaped by climatic, geographic, and ethnolinguistic factors. Two genes associated with seed size and shattering showed signatures of selection. Most known domestication genes from other cereal models however have not experienced strong selection in fonio, providing direct targets to rapidly improve this crop for agriculture in hot and dry environments.

## Introduction

Humanity faces the unprecedented challenge of having to sustainably produce healthy food for 9-10 billion people by 2050 in a context of climate change. A more efficient use of plant diversity and genetic resources in breeding has been recognized as a key priority to diversify and transform agriculture^1-3^. The Food and Agriculture Organization of the United Nations (FAO) stated that arid and semi-arid regions are the most vulnerable environments to increasing uncertainties in regional and global food production^4^. In most countries of Africa and the Middle East, agricultural productivity will decline in the near future^4^, because of climate change, land degradation, and groundwater depletion^5^. Agricultural selection, from the early steps of domestication to modern-day crop breeding, has resulted in a marked decrease in agrobiodiversity^6,7^. Today, three cereal crops alone, bread wheat (*Triticum aestivum*), maize (*Zea mays*), and rice (*Oryza sativa*) account for more than half of the globally consumed calories^8^.

Many of today’s major cereal crops, including rice and maize, originated in relatively humid tropical and sub-tropical regions^9,10^. Although plant breeding has adapted the major cereal crops to a wide range of climates and cultivation practices, there is limited genetic diversity within these few plant species for cultivation in the most extreme environments. On the other hand, crop wild relatives and orphan crops are often adapted to extreme environments and their utility to unlock marginal lands for agriculture has recently regained interest^2,6,11-14^. Current technological advances in genomics and genome editing provide an opportunity to rapidly domesticate wild relatives and to improve orphan crops^15,16^. *De novo* domestication of wild species or rapid improvement of semi-domesticated crops can be achieved in less than a decade by targeting a few key genes^6^.

White fonio (*Digitaria exilis* (Kippist) Stapf) (Fig. 1) is an indigenous African millet species with a great potential for agriculture in marginal environments^17,18^. Fonio is cultivated under a large range of environmental conditions, from a tropical monsoon climate in western Guinea to a hot, arid desert climate (BWh) in the Sahel zone. Some extra-early maturing fonio varieties produce mature grains in only 70-90 days^19^, which makes fonio one of the fastest maturing cereals. Because of its quick maturation, fonio is often grown to avoid food shortage during the lean season (period before main harvest), which is why fonio is also referred to as ‘hungry rice’. In addition, fonio is drought tolerant and adapted to nutrient-poor, sandy soils^20^. Despite its local importance, fonio shows many unfavorable characteristics that resemble undomesticated plants: seed shattering, lodging, and lower yields than other cereals^18^ (Supplementary Fig. 1). In the past few years, fonio has gained in popularity inside and outside of West Africa because of its nutritional qualities.

**Figure 1.**
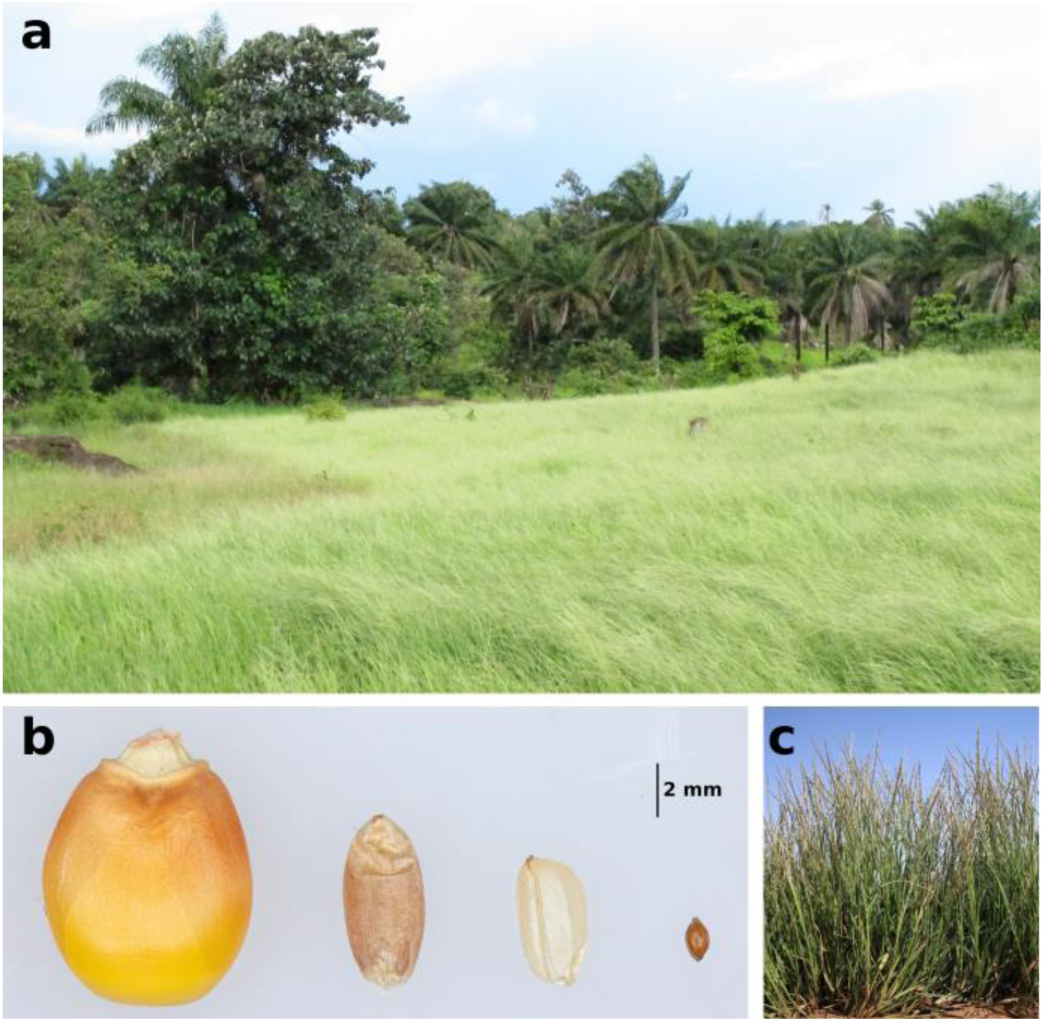
Phenotype of fonio (*Digitaria exilis*). **a**, Field of cultivated fonio in Guinea. **b**, Grains of maize, wheat, rice, and fonio (from left to right). **c**, Plants of the fonio accession CM05836.

Here, we present the establishment of a comprehensive set of genomic resources for fonio, which constitutes the first step towards harnessing the potential of this cereal crop for agriculture in harsh environments. These resources include the generation of a high-quality, chromosome-scale reference assembly and the deep re-sequencing of a diversity panel that includes wild and cultivated accessions covering a wide geographic range.

## Results

### Chromosome-scale fonio reference genome assembly

Fonio is a tetraploid species (2n = 4x = 36)^21^ with a high degree of selfing^17^. To build a *D. exilis* reference assembly, we chose an accession from one of the driest regions of fonio cultivation, CM05836 from the Mopti region in Mali. The size of the CM05836 genome was estimated to be 893 Mb/1C by flow cytometry (Supplementary Fig. 2 and 3), which is in line with previous reports^21^. The CM05836 genome was sequenced and assembled using deep sequencing of multiple short-read libraries (Supplementary Table 1), including Illumina paired-end (321-fold coverage), mate-pair (241-fold coverage) and linked-read (10x Genomics, 84-fold coverage) sequencing. The raw reads were assembled and scaffolded with the software package DeNovoMAGIC3 (NRGene), which has recently been used to assemble various high-quality plant genomes^22-24^. Integration of Hi-C reads (122-fold coverage, Supplementary Table 2) and a Bionano optical map (Supplementary Table 3) resulted in a chromosome-scale assembly with a total length of 716,471,022 bp, of which ∼91.5% (655,723,161 bp) were assembled in 18 pseudomolecules. A total of 60.75 Mb were unanchored (Table 1). Of 1,440 Embryophyta single copy core genes (BUSCO version 3.0.2), 96.1% were recovered in the CM05836 assembly, 2.9% were missing and 1% was fragmented. As no genetic *D. exilis* map is available, we used chromosome painting to further assess the quality of the CM05836 assembly. Pools of short oligonucleotides covering each one of the 18 pseudomolecules were designed based on the CM05836 assembly, fluorescently labelled, and hybridized to mitotic metaphase chromosome spreads of CM05836^25^. Each of the 18 libraries specifically hybridized to only one chromosome pair, confirming that our assembly unambiguously distinguished homoeologous chromosomes (Fig. 2a, Supplementary Fig. 4). Centromeric regions contained a tandem repeat with a 314 bp long unit, which was found in all fonio chromosomes (Supplementary Fig. 5). We also re-assembled all the data with the open-source TRITEX pipeline^26^ and the two assemblies showed a high degree of collinearity (Supplementary Fig. 6).

**Table 1.**
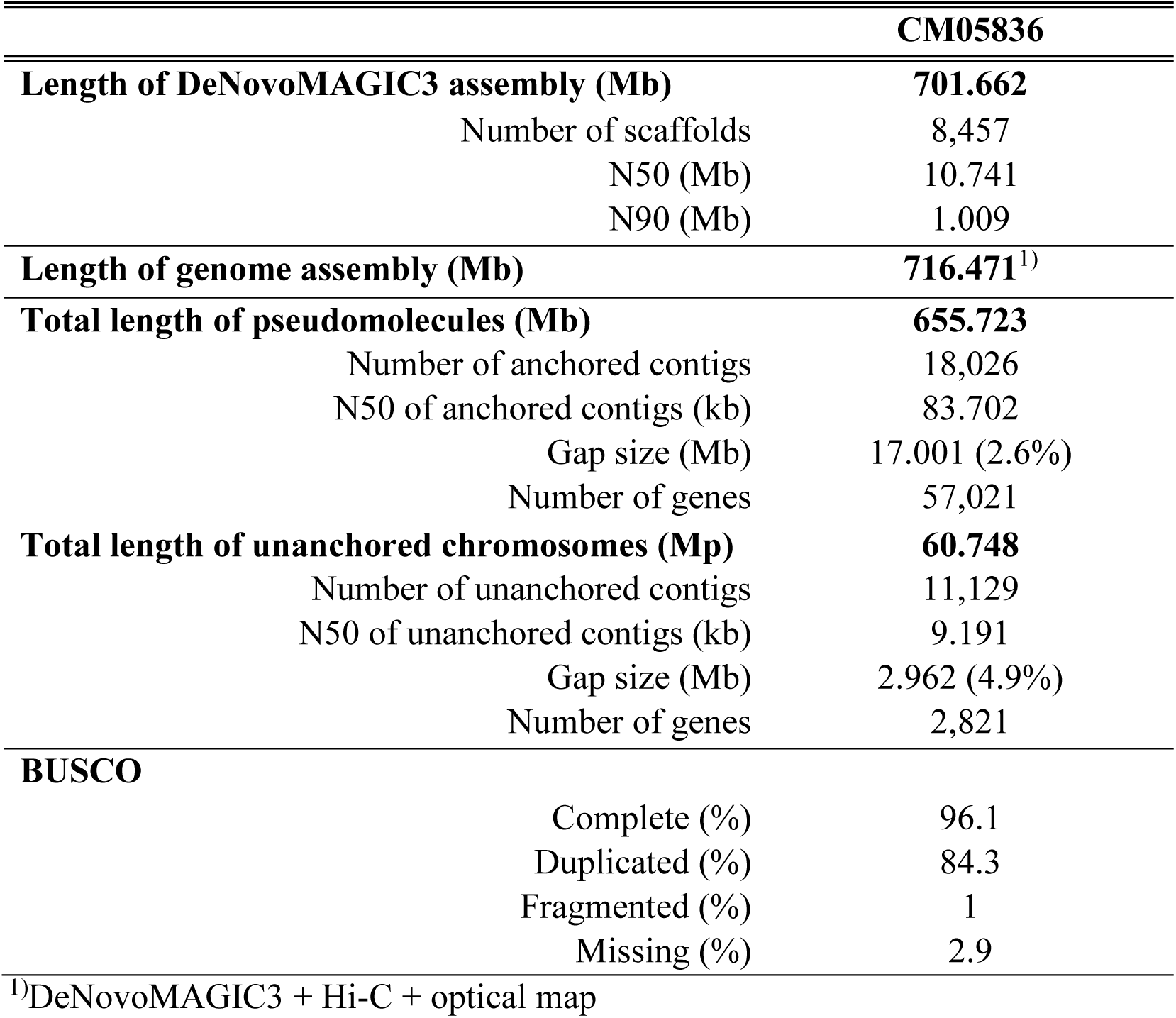
Statistics of the fonio genome assembly and annotation.

**Figure 2.**
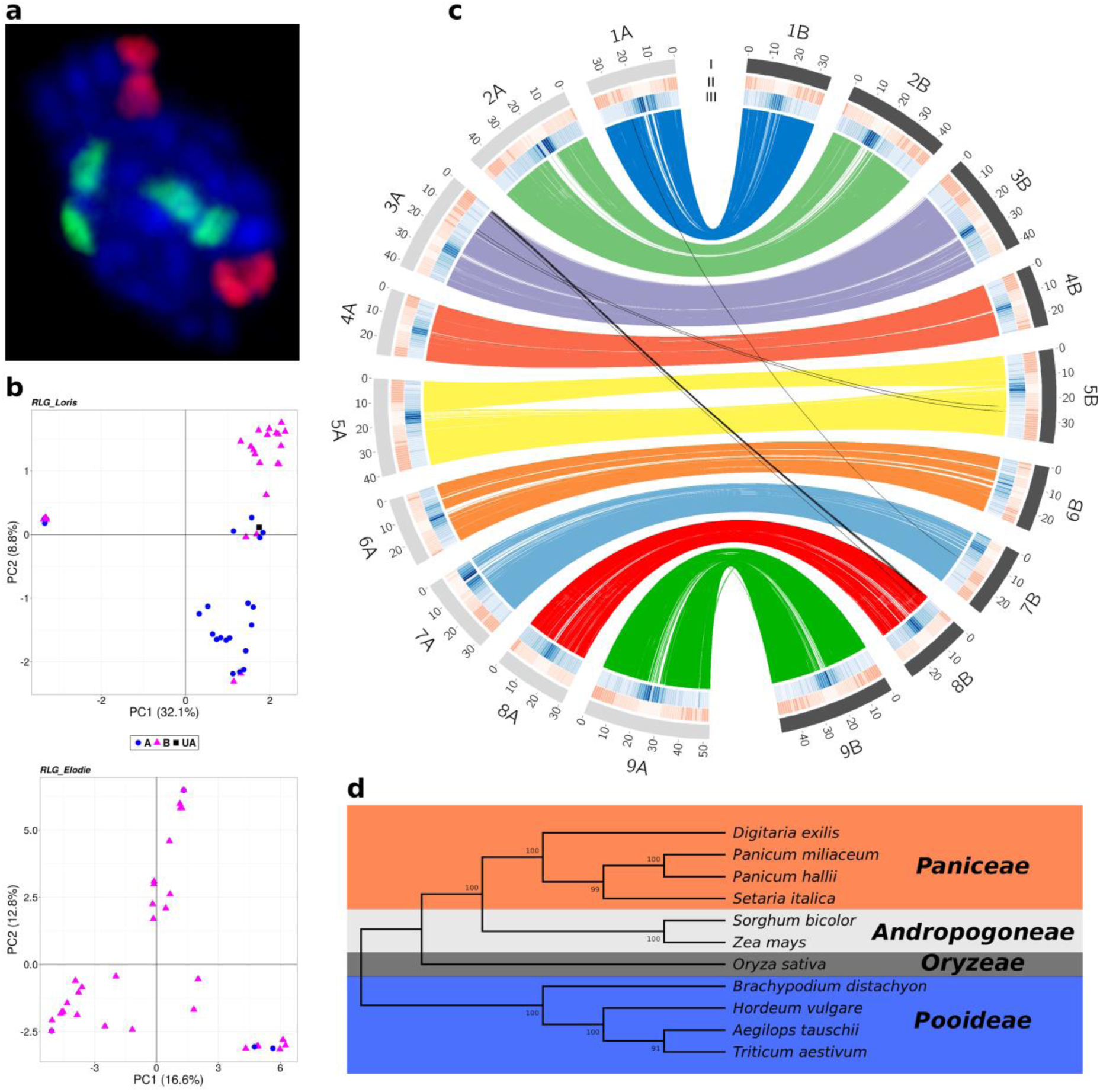
Fonio genome features. **a**, Representative example of oligo painting FISH on mitotic metaphase chromosomes. Shown are probes designed from pseudomolecules 9A (red) and 9B (green) of the CM05836 assembly. **b**, Principal Component Analysis (PCA) of the transposable element cluster RLG_Loris (upper panel) that allowed discrimination of the two sub-genomes. Blue dots represent elements found on the A sub-genome; pink triangles represent elements from the B sub-genome; black squares represent elements present on chromosome unanchored. PCA of the transposable element cluster RLG_Elodie (lower panel) that was specific to the B sub-genome. **c**, Synteny and distribution of genome features. (I) Number and length of the pseudomolecules. The grey and black colours represent the two sub-genomes. (II, III) Density of genes and repeats along each pseudomolecule, respectively. Lines in the inner circle represent the homoeologous relationships. **d**, Maximum likehood tree of eleven *Poaceae* species based on 30 orthologous gene groups. Topologies are supported by 1,000 bootstrap replicates. Colors indicate the different clades.

We compared the fonio pseudomolecule structure to foxtail millet (*Setaria italica*; 2n = 2x = 18), a diploid relative with a fully sequenced genome^27^. The fonio genome shows a syntenic relationship with the genome of foxtail millet, with two homoeologous sets of nine fonio chromosomes (Supplementary Fig. 7). Without a clear diploid ancestor, we could not directly disentangle the two sub-genomes^28^. We thus used a genetic structure approach based on full-length long terminal repeat retrotransposons (fl-LTR-RT). A total of eleven fl-LTR-RT families with more than 30 elements were identified and defined as a ‘populations’, allowing us to apply genetic structure analyses that are often used in population genomics^29^. We searched for fl-LTR-RT clusters that only appeared in one of the two homoeologous sub-genomes. Out of the eleven fl-LTR-RT populations analyzed, two allowed us to discriminate the sub-genomes (Fig. 2b, Supplementary Fig. 8). The two families belong to the Gypsy superfamily and dating of insertion time was estimated to be ∼1.56 million years ago (MYA) (57 elements, 0.06-4.67 MYA) and ∼1.14 MYA (36 elements, 0.39-1.96 MYA), respectively. The two putative sub-genomes were designated A and B and chromosome numbers were assigned based on the synteny with foxtail millet (Supplementary Fig. 7).

Gene annotation was performed using the MAKER pipeline (version 3.01.02) with 34.1% of the fonio genome masked as repetitive. Transcript sequences of CM05836 from flag leaves, grains, panicles, and whole above-ground seedlings (Supplementary Table 4) in combination with protein sequences of publicly available plant genomes were used to annotate the CM05836 assembly. This resulted in the annotation of 59,844 protein-coding genes (57,023 on 18 pseudomolecules and 2,821 on unanchored chromosome) with an average length of 2.5 kb and an average exon number of 4.6. The analysis of the four CM05836 RNA-seq samples showed that 44,542 protein coding genes (74.3%) were expressed (>0.5 transcripts per million), which is comparable to the annotation of the bread wheat genome (Supplementary Table 5)^30,31^.

### Synteny with other cereals

Whole genome comparative analyses of the CM05836 genome with other grass species were consistent with the previously established phylogenetic relationships of fonio^20^. A comparison of the CM05836 A and B sub-genomes identified a set of 16,514 homoeologous gene pairs that fulfilled the criteria for evolutionary analyses (Fig. 2c and Supplementary data 1). The estimation of synonymous substitution rates (Ks) among homoeologous gene pairs revealed a divergence time of the two sub-genomes of roughly 3 MYA (Supplementary Fig. 9). These results indicate that *D. exilis* is a recent allotetraploid species. A Ks distribution using orthologous genes revealed that fonio diverged from the other members of the *Paniceae* tribe (broomcorn millet (*Panicum miliaceum*), Hall’s panicgrass (*P. hallii*), and foxtail millet (*S. italica*)) between 14.6 and 16.9 MYA, and from the *Andropogoneae* tribe (sorghum (*Sorghum bicolor*) and maize (*Z. mays*)) between 21.5 and 26.9 MYA. Bread wheat (*T. aestivum*), goatgrass (*Aegilops tauschii*), barley (*Hordeum vulgare*), rice (*O. sativa*), and purple false brome (*Brachypodium distachyon*) showed a divergence time of 35.3 to 40 MYA (Fig. 2d).

The hybridization of two genomes can result in genome instability, extensive gene loss, and reshuffling. As a consequence, one sub-genome may evolve dominancy over the other sub-genome^23,30,32-34^. Using foxtail millet as a reference, the fonio A and B sub-genomes showed similar numbers of orthologous genes: 14,235 and 14,153 respectively. Out of these, 12,541 were retained as duplicates while 1,694 and 1,612 were specific to the A and B sub-genome, respectively. The absence of sub-genome dominance was also supported by similar gene expression levels between homoeologous pairs of genes (Supplementary Fig. 10 and Supplementary data 2).

### Evolutionary history of fonio and its wild relative

To get an overview of the diversity and evolution of fonio, we selected 166 *D. exilis* accessions originating from Guinea, Mali, Benin, Togo, Burkina Faso, Ghana, and Niger and 17 accessions of the proposed wild tetraploid fonio progenitor *D. longiflora*^35^ from Cameroon, Nigeria, Guinea, Chad, Soudan, Kenya, Gabon, and Congo for whole-genome re-sequencing. The selection was done from a collection of 641 georeferenced *D. exilis* accessions^36^ with the aim of maximizing diversity based on bioclimatic data and geographic origin. We obtained short-read sequences with an average of 45-fold coverage for *D. exili*s (range 36 - 61-fold) and 20-fold coverage for *D. longiflora* (range 10 - 28-fold) (Supplementary Table 6). The average mapping rates of the raw reads to the CM05836 reference assembly were 85% and 68% for *D. exilis* and *D. longiflora*, respectively, with most accessions showing a mapping rate of >80% (Supplementary Table 7).

After filtering, 11,046,501 high quality bi-allelic single nucleotide polymorphisms (SNPs) were retained. Nine *D. exilis* and three *D. longiflora* accessions were discarded based on the amount of missing data. The SNPs were evenly distributed across the 18 *D. exilis* chromosomes, with a tendency toward a lower SNP density at the chromosome ends (Supplementary Fig. 11). Most of the SNPs (84.3%) were located in intergenic regions, 9.5% in introns and 6.2% in exons. Of the exonic SNPs, 354,854 (51.6%) resulted in non-synonymous sequence changes, of which 6,727 disrupted the coding sequence (premature stop codon). The remaining 333,296 (48.4%) exonic SNPs represented synonymous variants. Forty-four percent of the total SNPs (4,901,160 SNPs) were rare variants with a minor allele frequency of less than 0.01 (Supplementary Fig. 12). The mean nucleotide diversity (*π*) was 6.19 × 10^−4^ and 3.68 × 10^−3^ for *D. exilis* and *D. longiflora*, respectively. Genome-wide linkage disequilibrium (LD) analyses revealed a faster LD decay in *D. longiflora* (r^2^ ∼ 0.16 at 70 kb) compared to *D. exilis* (r^2^ ∼ 0.20 at 70 kb) (Supplementary Fig. 13).

A PCA showed a clear genetic differentiation between cultivated *D. exilis* and wild *D. longiflora*. The *D. exilis* accessions clustered closely together, while the *D. longiflora* accessions split into three groups (Fig. 3a). The *D. longiflora* group that showed the greatest genetic distance from *D. exilis* contained three accessions originating from Central (Cameroon) and East Africa (South Sudan and Kenya). The geographical projection of the first principal component (which separated wild accessions from the cultivated accessions) revealed that the *D. exilis* accessions genetically closest to *D. longiflora* originated from southern Togo and the west of Guinea (Supplementary Fig. 14a).

**Figure 3.**
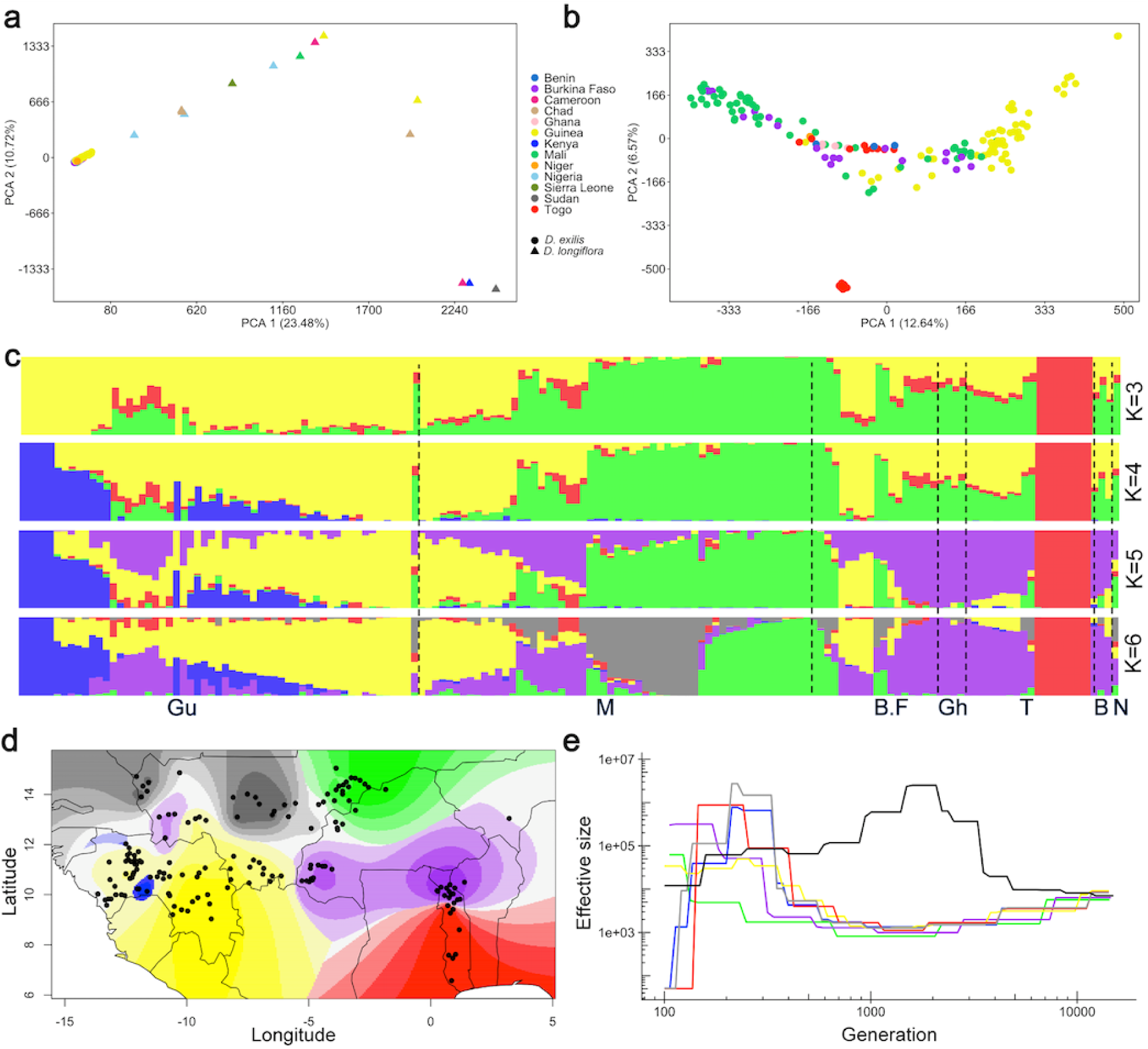
Genetic diversity and structure of *D. exilis* and *D. longiflora* diversity panel. **a**, Principal component analysis (PCA) of 157 *D. exilis* and 14 *D. longiflora* accessions using whole-genome single nucleotide polymorphisms (SNPs). *D. exilis* samples (circles), *D. longiflora* (triangles). **b**, PCA of *D. exilis* accessions alone. Colors indicate the country of origin. **c**, Population structure (from *K*=3 to *K*=6) of *D. exilis* accessions estimated with sNMF. Each bar represents an accession and the bars are filled by colors representing the likelihood of membership to each ancestry. Accessions are ordered from west to east; Guinea (Gu), Mali (M), Burkina Faso (B.F), Ghana (Gh), Togo (T), Benin (B), and Niger (N). **d**, Geographic distribution of ancestry proportions of *D. exilis* accessions obtained from the structure analysis. The colors represent the maximal local contribution of an ancestry. Black dots represent the coordinates of *D. exilis* accessions. **e**, Effective population size history of *D. exilis* groups based on *K*=6 and *D. longiflora* (in black).

A PCA with *D. exilis* accessions alone revealed three main clusters. The second principal component separated eight accessions from southern Togo from the remaining accessions (Fig. 3b). In addition, five accessions from Guinea formed a distinct genetic group in the PCA. The remaining accessions were spread along the first axis of the PCA, mainly revealing a grouping by geographic location (Fig. 3b). The genetic clustering was confirmed by genetic structure analyses (Fig. 3c and d). The cross-validation error decreased with increasing *K* and reached a plateau starting from *K*=6 (Supplementary Fig. 14b). At *K*=3, the eight South Togo accessions formed a distinct homogenous population. At *K*=4, the five accessions from Guinea were separated. With increasing *K*, the admixture plot provided some evidence that natural (climate and geography) and human (ethnicity and language) factors had an effect on shaping the genetic population structure of fonio (Fig. 3c and d, Supplementary Fig. 14c). We observed a significant correlation (Pearson’s correlation; p < 0.05) between the genetic population structure (first principal component of PCA) and climate (i.e. mean temperature and precipitation of the wettest quarters) as well as geography (i.e. latitude, longitude and altitude) (Fig. 3b, Supplementary Fig. 15, Supplementary Table 8). A significant correlation was also observed for ethnic and linguistic groups (fisher test: p-value = 0.0005, mantel test: p-value = 0.001). The effect of ethnic groups remained significant (p = 0.045) even when we controlled for geographic and climatic factors (ANCOVA) (Supplementary Table 9). Unadmixed populations were mainly found at the geographic extremes of the fonio cultivation area in the north and south, whereas the accessions from the central regions of West Africa tended to show a higher degree of admixture (Supplementary Fig. 16a).

Plotting the spatial distribution of private SNPs (i.e., SNPs present only once in a single genotype) revealed a hotspot of rare alleles in Togo, Niger, the western part of Guinea, and southern Mali (Supplementary Fig. 16b and c). Rare allele diversity was lower in the eastern part of Guinea, and in southwest Mali.

Inference of the *D. exilis* effective population size revealed a decline that started more than 10,000 years ago and reached a minimum between 2,000 and 1,000 years ago (Fig. 3e). Then, a steep increase of the effective population size occurred to a level that was approximately 100-fold higher compared to the bottleneck.

### Genomic footprints of selection and domestication

We used three complementary approaches to detect genomic regions under selection: (*i*) a composite likelihood-ratio (CLR) test based on site frequency spectrum (SFS)^37^, (*ii*) the nucleotide diversity (π) ratios between *D. exilis* and *D. longiflora* over sliding genomic windows, and (*iii*) the genetic differentiation based on *F*_*ST*_ calculations, again computed over sliding windows. With the CLR test, 78 regions were identified as candidates for signatures of selection. The genetic diversity ratios and *F*_*ST*_ calculations revealed 311 and 208 regions, respectively (Fig. 4, Supplementary Data 3). We then searched for the presence of orthologs of known domestication genes in the regions under selection. The most striking candidate was one of the two orthologs of the rice grain size *GS5* gene^38^ (Dexi3A01G0012320 referred to as *DeGS5-3A*) that was detected by genetic diversity ratio and *F*_*ST*_ based calculations (Fig. 4). *GS5* regulates grain width and weight in rice. *D. exilis* showed a dramatic loss of genetic diversity at the *DeGS5-3A* gene (Fig. 5a and b). Domestication of fonio is associated with wider grains of *D. exilis* compared to grains of *D. longiflora* (Fig. 5c). The region of the *GS5* ortholog on chromosome 3B (*DeGS5-3B*) was not identified as being under selection and showed higher levels of nucleotide diversity than the *DeGS5-3A* region (Fig. 5a and b). Only the *DeGS5-3A* but not the *DeGS5-3B* transcript was detected in the *D. exilis* RNA-Seq data from grains. This is in agreement with observations made in rice^38^, where a dominant mutation resulting in increased *GS5* transcript levels affects grain size.

**Figure 4.**
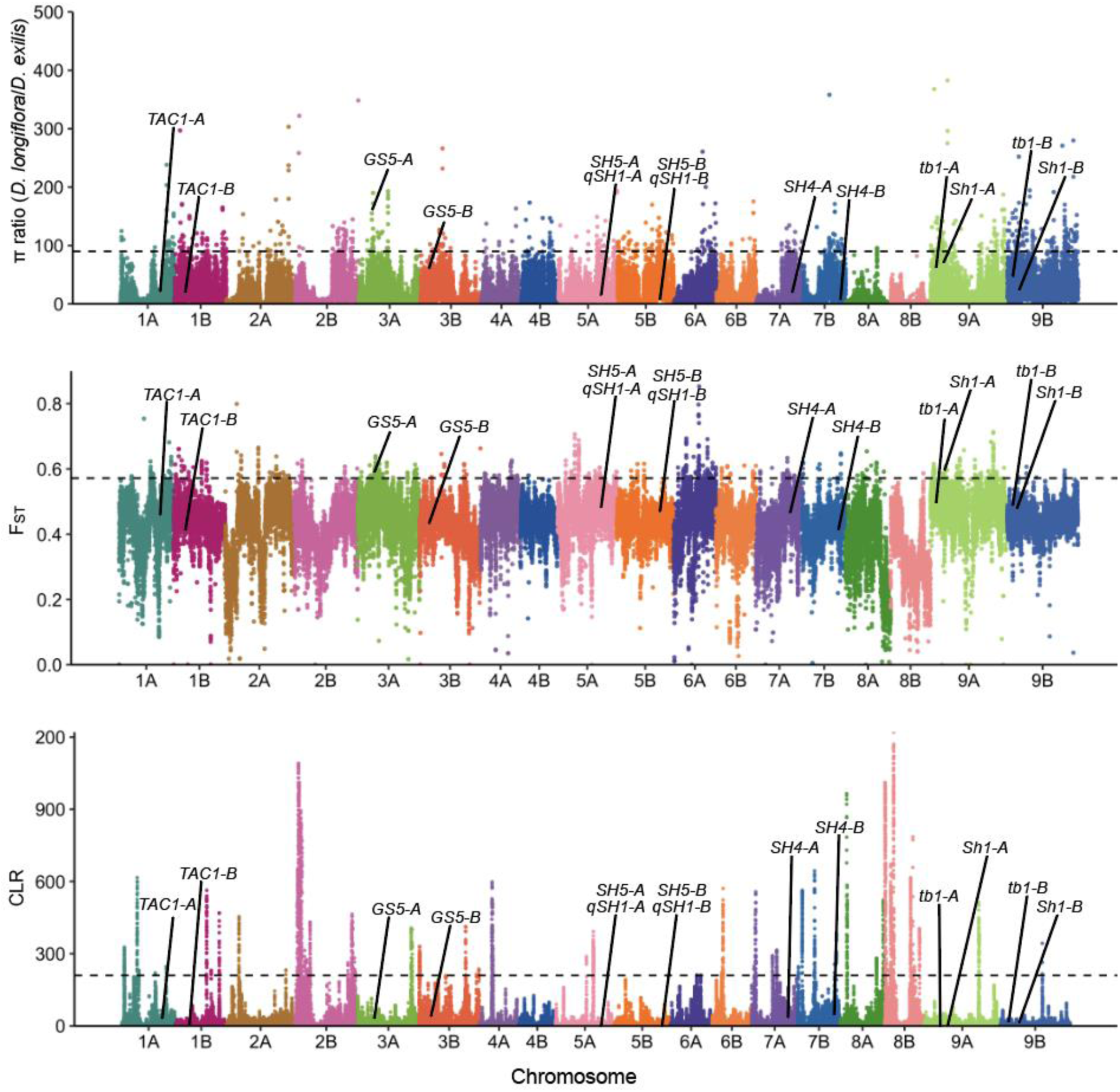
Detection of selection in fonio. Manhattan plots showing detection of selection along the genome based on π ratio, F_ST_ and SweeD (from top to bottom). The location of orthologous genes of major seed shattering and plant architecture genes are indicated in the Manhattan plots. The black dashed lines indicate the 1% threshold. Some extreme outliers in the π ratio plot are not shown.

**Figure 5.**
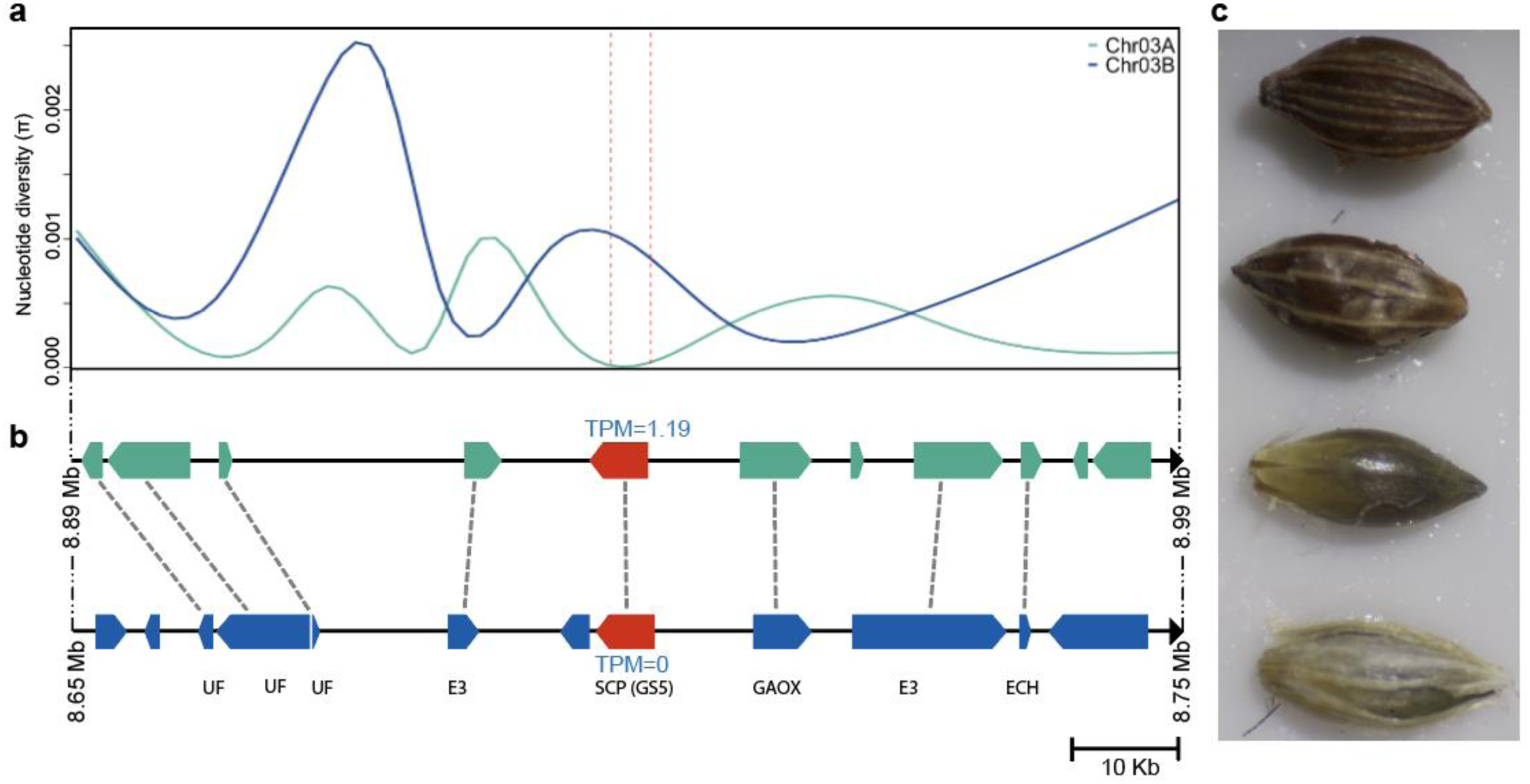
Selective sweep at the *GS5* locus in fonio. **a**, Smoothed representation of nucleotide diversity (π) in *D. exilis* in a 100 kb window surrounding the ortholog of the rice *GS5* gene. The green curve shows π around *DeGS5-3A* on chromosome 3A and the blue curve π around *DeGS5-3B* on chromosome 3B. The dashed vertical red lines represent the location of the *GS5* genes. The nucleotide diversity was calculated in overlapping 100 bp windows every 25 bp. **b**, Schematic representation of the annotated genes in the *GS5* regions and the orthologous gene relationships between chromosomes 3A (green) and 3B (blue). *UF*, = protein of unknown function, *E3* = E3 ubiquitin ligase; *SCP* (*GS5*) = serine carboxypeptidase (Dexi3A01G0012320); *GAOX* = gibberellin 2-beta-dioxygenase; *ECH* = golgi apparatus membrane protein. *DeGS5-3A* and *DeGS5-3B* are indicated in red. The numbers in blue above and below the *GS5* orthologs show their respective expression in transcripts per million (TPM) in grain tissue. **c**, Two grains of *D. exilis* (top) and *D. longiflora* (bottom).

Another domestication gene that was detected in the selection scan was an ortholog of the sorghum *Shattering 1* (*Sh1*) gene^39^. *Sh1* encodes a YABBY transcription factor and the non-shattering phenotype in cultivated sorghum is associated with lower expression levels of *Sh1* (mutations in regulatory regions or introns) or truncated transcripts. Domesticated African rice (*O. glaberrima*) carries a 45 kb deletion at the orthologous *OsSh1* locus compared to its wild relative *O. barthii*^40^. Around 37% of the fonio accessions had a 60 kb deletion similar to *O. glaberrima* that eliminated the *Sh1* ortholog on chromosome 9A (*DeSh1-9A -* Dexi9A01G0015055) (Fig 6a). The homoeologous region including *DeSh1-9B* (Dexi9B01G0013485) on chromosome 9B was intact. Interestingly, accessions with the *DeSh1-9A* deletion showed a minor (7%) but significant reduction in seed shattering (Fig. 6b) compared to accessions with the intact *DeSh1-9A* gene. Accessions carrying this deletion were distributed across the whole range of fonio cultivation, which suggests that the deletion is ancient and might have been selected for in certain regions.

**Figure 6.**
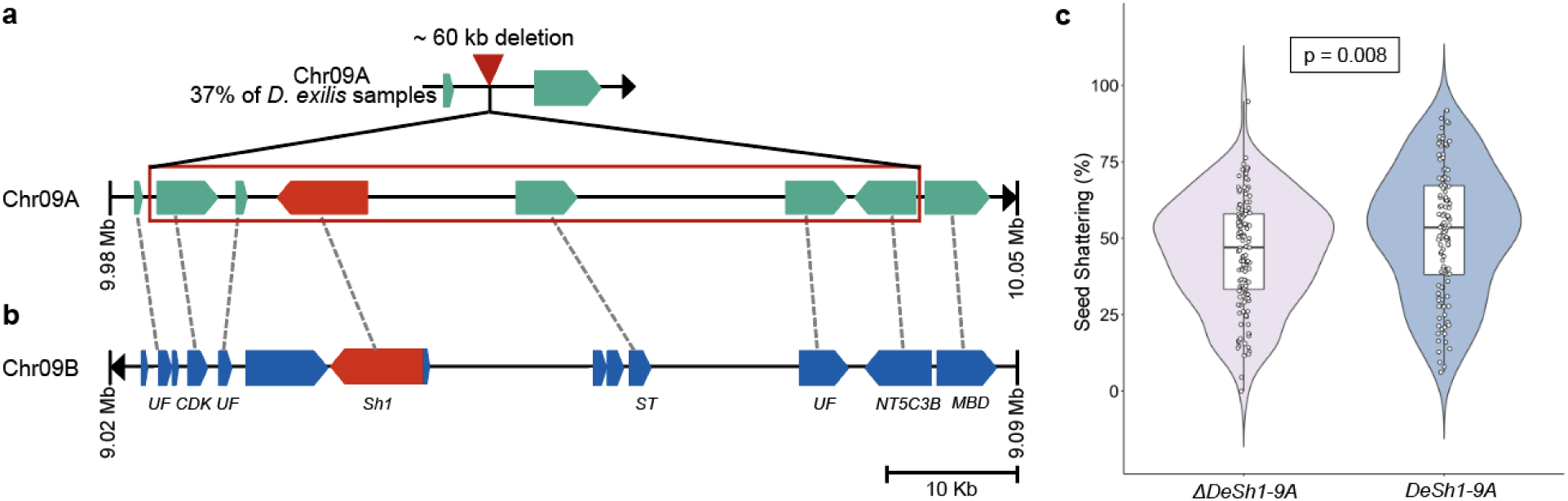
Selective sweep at the *Sh1* locus in fonio. **a**, Schematic representation of genes in the orthologous regions of the sorghum *Sh1* gene. The top most panel shows the region on fonio chromosome 9A with the 60 kb deletion as it is found in 37% of all *D. exilis* accessions. The lower panel shows the genes in accessions without the deletion. The *Sh1* ortholog *DeSh1-9A* is shown in red. **b**, The *Sh1* ortholog *DeSh1-9B* on chromosome 9B is present in all *D. exilis* accessions. The dashed lines represent orthologous relationships between the two sub-genomes. *UF* = protein of unknown function; *CDK* = cyclin-dependent kinase; *Sh1* = ortholog of sorghum *Shattering 1* (*Sh1*); *ST* = sulfate transporter; *NT5C3B* = 7-methylguanosine phosphate-specific 5’-nucleotidase; *MBD* = methyl-CpG-binding domain. **c**, Seed shattering in accessions carrying the *DeSh1-9A* deletion (*ΔDeSh1-9A*) compared to accessions with an intact *DeSh1-9A*. Mean seed shattering was 45% in *ΔDeSh1-9A* (± 0.18 s.d.) and 52% in *DeSh1-9A* (± 0.22 s.d.). (n = Three individual panicles of 43 Δ*DeSh1-9A* accessions and 39 *DeSh-9A* accessions, respectively).

## Discussion

Here, we established a set of genomic resources that allowed us to comprehensively assess the genetic variation found in fonio, a cereal crop that holds great promises for agriculture in marginal environments. The analysis of fl-LTR-RT revealed two sub-genome specific transposon clusters that experienced a peak of activity around 1.1 - 1.5 MYA, indicating that the two fonio sub-genomes hybridized after this period^41^. The Ks analysis estimated that the two sub-genomes diverged prior to these transposon bursts ∼ 3 MYA, suggesting that fonio is an allotetraploid species. The analysis of effective population size revealed a genetic bottleneck that was most likely associated with human cultivation and domestication. The large increase of effective size for fonio after a period of reduction was most probably due to the development and expansion of fonio cultivation. This expansion appears to be a recent event and occurred during the last millennium. The progression of the effective population size resembles the patterns observed for other domesticated crops, with a protracted period of cultivation followed by a marked bottleneck^42,43^. In the case of fonio, this potential bottleneck appears to be milder compared to other crops. It has been observed that the effective population size of the wild rice (*O. barthii*) from West Africa followed a trend similar to the cultivated African rice (*O. glaberrima*), which has been interpreted as a result of environmental degradation^42^ rather than human selection. In contrast, no signal of population bottleneck was observed for the proposed wild fonio ancestor *D. longiflora*, indicating that the bottleneck observed in fonio is associated with human cultivation. We also highlighted the strong impact of geographic, climatic and anthropogenic factors on shaping the genetic diversity of fonio. Even if fonio is not a dominant crop across West Africa, it benefits from cultural embedding and plays a key role in ritual systems in many African cultures^44^. For example, we observed a striking genetic differentiation between *D. exilis* accessions collected from northern and southern Togo. This can be attributed to both climatic and cultural differences. While the southern regions of Togo receive high annual rainfalls, the northern parts receive less than 1,000 mm annual rainfall and there is a prolonged dry season^45^. Adoukonou-Sagbadja et al.^45^ also noted that there is no seed exchange between farmers of the two regions because of cultural factors.

Despite a genetic bottleneck and the reduction in genetic diversity, fonio still shows many ‘wild’ characteristics such as residual seed shattering, lack of apical dominance, and lodging. We show that orthologs of most of the well-characterized domestication genes from other cereals were not under strong selection in fonio. Interestingly, an ortholog of the rice *GS5* gene (*DeGS5-3A*) was identified in a selective sweep. Dominant mutations in the *GS5* promoter region were associated with higher *GS5* transcript levels and wider and heavier grains in rice^38^. The *DeGS5-3A* gene showed a complete loss of diversity in the coding sequence in *D. exilis*, suggesting a strong artificial selection for larger grains. The hard selective sweep at the *DeGS5-3A* locus is in contrast to *DeSh1-9A*, which showed evidence for a partial selective sweep^46^. The non-shattering phenotype associated with the *Sh1* locus in sorghum is recessive^39^ and the deletion of a single *DeSh1* copy only resulted in a quantitative reduction of shattering that might have been selected for in some but not all regions. Whether the 60 kb deletion including *DeSh1-9A* represents standing genetic variation or arose after fonio domestication cannot be determined. The deletion was not identified in any of the 14 re-sequenced *D. longiflora* accessions. Targeting the *DeSh1-9B* locus on chromosome 9B in accessions that carry the *DeSh1-9A* deletion through mutagenesis or genome editing could produce a fonio cultivar with significantly reduced seed shattering, which would form a first important step towards a significant improvement of this crop.

## Acknowledgements

We are grateful to Samantha Bazan, Ablaye Ngom, Marie Piquet, Hélène Adam and Carole Gauron for their technical assistance in Herbarium sampling and photography. Cirad herbarium samples of *Digitaria longiflora* were provided by Samantha Bazan (ALF, http://publish.plantnet-project.org/). We thank Soukeye Conde for valuable insight, funded by the project Cultivar, reference ANR-16-IDEX-0006, through the Investissements d’avenir program (Labex Agro: ANR-10-LABX-0001-01). This work was supported by the CIRAD-UMR AGAP and IRD UMR DIADE HPC Data Center of the South Green Bioinformatics platform (http://www.southgreen.fr/) and the King Abdullah University of Science and Technology (KAUST). D.S., J.C., E.H. and J.D. were in part supported by the European Regional Development Fund OPVVV project “Plants as a tool for sustainable development” number CZ.02.1.01/0.0/0.0/16_019/0000827.

## Author contributions

M.A., A.B., Y.V. and S.G.K. designed research. N.K. and M.A. established the SNP calling pipeline. J.B., M.C. and L.Z. performed molecular analyses and constructed sequencing libraries. M.A., H.I.A., P.C., L.G., N.S. and T.W. performed bioinformatics analyses. M.A., H.I.A., P.C., Y.V., C.B., A.B. and S.G.K. interpret results. D.S., J.C., E.H. and J.D. performed flow-cytometry and chromosome painting. S.A., S.C. and H.B. constructed and analysed the CM05836 optical map. Y.P. and H.I.A carried out phenotypic data collection and analyses. J.J.W., M.G., N.A.K., C.L., S.C., S.V. and C.B. contributed biological materials. M.A., H.I.A. and S.G.K. wrote the paper with substantial inputs from P.C., Y.V. and A.B. All authors have read and approved the manuscript.

## Additional information

Supplementary information is available for this paper

Correspondence and requests for materials should be addressed to A.B. and S.G.K.

## Methods

### Plant material

The plant material used in this study comprised a collection of 641 *D. exilis* accessions maintained at the French National Research Institute for Development (IRD), Montpellier, France^36,47^. The collection was established from 1977 to 1988. For genome re-sequencing, a panel of 166 accessions (Supplementary Table 8) was selected based on geographic and bioclimatic data. One plant per accession was chosen for DNA extraction and sequencing. Inflorescences of sequenced *D. exilis* plants were covered with bags to prevent outcrossing. Grains of sequenced plants were collected and kept for further analyses. In addition, 17 accessions of *D. longiflora* were collected from specimens stored at the National Museum of Natural History of Paris (Paris, France), IFAN Ch. A. Diop (Dakar, Senegal), National Herbarium of The Netherlands (Leiden, Netherlands), and Cirad (Montpellier, France) (Supplementary Table 10)

### Flow cytometry for genome size estimation

The amount of nuclear DNA in fonio was measured by flow-cytometry^48^. In brief, six different individual plants representing accession CM05836 were measured three times on three different days using a CyFlow^®^ Space flow cytometer (Sysmex Partec GmbH, Görlitz, Germany) equipped with a 532 nm green laser. Soybean (*Glycine max* cultivar Polanka; 2C = 2.5 pg) was used as an internal reference standard. 2C DNA contents (in pg) were calculated from the mean G1 peak (interphase nuclei in G1 phase) positions by applying the formula:

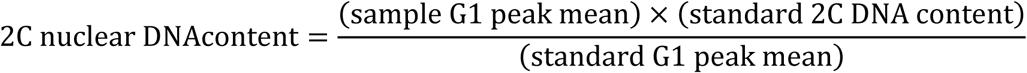

DNA contents in pg were converted to genome sizes in bp using the conversion factor 1 pg DNA = 0.978 Gb^49^.

### Genome sequencing and assembly

High molecular weight (HMW) genomic DNA was isolated from a single CM05836 plant with a dark-treatment of 48 hours before harvesting tissue from young leaves. Two paired-end and three mate-pair libraries were constructed with different insert sizes ranging from 450 bp to 10 kb. The 450 bp paired-end library was sequenced on a Illumina Hi-Seq 2500 instrument. The other libraries were sequenced on a Illumina NovaSeq 6000 instrument. In addition, one 10x linked read library was constructed and sequenced with a Illumina NovaSeq 6000 instrument, yielding 75-fold coverage. A *de novo* whole genome assembly (WGA) was performed with the DeNovoMAGIC3 software (NRGene).

For the super-scaffolding, one Hi-C library was generated using the Dovetail™ Hi-C Library Preparation Kit and sequenced on one lane of a Illumina HiSeq 4000 instrument, yielding a ∼128-fold coverage. The assembly to Hi-C super-scaffolds was performed with the HiRise™ software. This generated 18 large super-scaffolds with a size between 25.5 Mb and 49.1 Mb (total size = 642.8 Mb with N50 = 39.9 Mb), while the 19^th^ longest super-scaffold was only 1.5 Mb in size, indicating that the 18 longest super-scaffolds correspond to the 18 *D. exilis* chromosomes.

To generate a BioNano optical map, ultra HMW DNA was isolated using the QIAGEN Genomic-tip 500/G kit (Cat No./ID: 10262) from plants grown in a greenhouse with a dark treatment applied during 48 h prior to collecting leaf tissue. Labeling and staining of the HMW DNA were performed according to the Bionano Prep Direct Label and Stain (DLS) protocol (30206 - Bionano Genomics) and then loaded on one Saphyr chip. The optical map was generated using the Bionano Genomics Saphyr System according the Saphyr System User Guide (30247 - Bionano Genomics). A total of 1,114.6 Gb of data were generated, but only 210.6 Gb of data corresponding to molecules with a size larger than 150 kb were retained. Hybrid scaffolding between the WGA and the optical maps was done with the hybridScaffold pipeline (with the Bionano Access default parameters). In total, 39 hybrid scaffolds ranging from 550 kb to 40.9 Mb (total length 657.3 Mb with N50 = 22.6 Mb) were obtained (Supplementary Table 3).

To construct the final chromosome-scale assembly, we integrated the Hi-C super-scaffolds and the hybrid scaffold manually. The 39 hybrid scaffolds generated with the optical map were aligned to the 18 Hi-C super-scaffolds produced by HiRise (Supplementary Table 11). The hybrid scaffolds detected two chimeric DeNovoMAGIC3 scafflods that most likely arose from wrong connections in the putative centromeres. The final chromosome-level assembly was constructed as follows: (i) We used the hybrid assembly to break the two chimeric DeNovoMAGIC3 scaffolds, (ii) merged and oriented the hybrid scaffolds using the Hi-C super scaffolds, and (iii) merged the remaining contigs and scaffolds into an unanchored chromosome.

A second genome assembly was produced using the TRITEX pipeline^26^. Briefly, the initial step using Illumina sequencing data (pre-processing of paired-end and mate-pair reads, unitig assembly, scaffolding and gap closing) was done according the instructions of the TRITEX pipeline (https://tritexassembly.bitbucket.io/). Integration of 10x and Hi-C reads was done differently. For the 10x scaffolding, we used tigmint v1.1.2^50^ to detect and cut sequences at positions with few spanning molecules, arks v1.0.3^51^ to generate graphs of scaffolds with connection evidence, and LINKS^52^ for a second step of scaffolding. For the Hi-C data, we used BWA^53^ to map the reads against the previous scaffolds and juicer tools v1.5^54^ for the super-scaffolding.

### Preparation of FISH probes and cytogenetic analyses

Libraries of 45 bp long oligomers specific for each fonio pseudomolecule were designed using the Chorus software (https://github.com/forrestzhang/Chorus) following the criteria of Han, et al.^25^. The number of oligomers per pseudomolecule was adjusted to ensure uniform fluorescent signals along the entire chromosomes after fluorescence *in situ* hybridization (FISH). In total, 310,484 oligomers were selected (between 12,647 and 20,000 oligomers per library) and were synthesized by Arbor Biosciences (Ann Arbor, MI, USA). To prepare chromosome painting probes for FISH, the oligomer libraries were labeled directly using 6-FAM or CY3, or indirectly using digoxigenin or biotin. A tandem repeat CL10 with a 314 bp long repetitive unit (155 bp long subunit) was identified after the analysis of fonio DNA repeats with the RepeatExplorer software^55^. A 20 bp oligomer was designed based on the CL10 sequence using the Primer3 software^56^ labeled by CY3 and used for FISH.

Mitotic metaphase chromosome spreads were prepared from actively growing roots (∼1 cm long) of fonio accession CM05836 according to Šimoníková et al.^57^. FISH was done with the chromosome painting probes and a probe for the CL10 tandem repeat following the protocol of Šimoníková et al.^57^. Digoxigenin- and biotin-labeled probes were detected using anti-digoxigenin-FITC (Roche Applied Science) and streptavidin-Cy3 (ThermoFisher Scientific/Invitrogen), respectively. The preparations were mounted in Vectashield with DAPI (Vector laboratories, Ltd., Peterborough, UK) to counterstain the chromosomes and the slides were examined with an Axio Imager Z.2 Zeiss microscope (Zeiss, Oberkochen, Germany) equipped with a Cool Cube 1 camera (Metasystems, Altlussheim, Germany) and appropriate optical filters. The capture of fluorescence signals and measure of chromosome lengths were done using the ISIS software 5.4.7 (Metasystems) and final image adjustment was done in Adobe Photoshop CS5.

### Analysis of full-length long terminal repeat retrotransposons (fl-LTR-RT)

fl-LTR-RT were identified using both LTRharvest^58^ (-minlenltr 100 -maxlenltr 40000 -mintsd 4 - maxtsd 6 -motif TGCA -motifmis 1 -similar 85 -vic 10 -seed 20 -seqids yes) and LTR_finder^59^ (-D 40000 -d 100 -L 9000 -l 50 -p 20 -C -M 0.9). Then, the candidate LTRs were filtered using LTR_retriever^60^.

fl-LTRs were classified into families using a clustering approach using MeShClust2^61^ with default parameters and manually curated with dotter^62^ to discard wrongly assigned LTRs. Only LTR-RT families with at least 30 intact copies were considered in genetics analyses, where different families represented ‘populations’ and each fl-LTR-RT copy of a family was considered as an individual. A multiple sequence alignment was performed within each family using Clustal Omega^63^ and the polymorphic sites were identified and converted into Variant Call Format (VCF) using msa2vcf.jar (https://github.com/lindenb/jvarkit). A principal component analysis (PCA) was performed for each fl-LTR-RT family using the R packages vcfR v.1.8.0^64^ and adegenet v.2.1.1^65^. Results were visualized with ggplot2^66^.

The approximate insertion dates of the fl-LTR-RTs were calculated using the evolutionary distance between two LTR sequences with the formula T=K/2μ, where K is the divergence rate approximated by percent identity and μ is the neutral mutation rate. The mutation rate used was μ=1.3 × 10^−8^ mutations per bp per year^67^.

### Gene annotation

A combination of homology-based and *de novo* approaches was used for repeat annotation using the RepeatModeler software (http://www.repeatmasker.org/RepeatModeler/) and the RepeatExplorer software^55^. The results of RepeatModeler, RepeatExplorer and LTR_retriever were merged into a comprehensive *de novo* repeat library using USEARCH v.11^68^ with default parameters (-id 0.9). The final repeats were classified using the RepeatClassifier module with the NCBI engine and then used to mask the CM05836 assembly.

The protein coding genes were predicted on the masked genome using MAKER (version 3.01.02)^69^. Transcriptome reads from flag leaf, grain, panicle and above-ground seedlings were filtered for ribosomal RNA using SortMeRNA v2.1^70^ and for adaptor sequences, quality and length using trimmomatic v0.38^71^. Filtered reads were mapped to the reference sequence using STAR v.2.7.0d^72^ and the alignments were assembled with StringTie v.1.3.5^73^ to further be used as transcript evidence. Protein sequences from *Arabidopsis thaliana*^74^, *Setaria italica*^27^, *Sorghum bicolor*^*75*^, *Zea mays*^76^ and *Oryza sativa*^77^ were compared with BLASTX^78^ to the masked pseudomolecules of CM05836 and alignments were filtered with Exonerate v.2.2.0^79^ to search for the accurately spliced alignments. A *de novo* prediction was performed using *ab initio* softwares GeneMark-ES v.3.54^80^, SNAP v.2006-07-28^81^ and Augustus v.2.5.5^82^. Three successive iterations of the pipeline MAKER were performed in order to improve the inference of the gene models and to integrate the final consensus genes. Finally, the putative gene functions were assigned using the UniProt/SwissProt database (Release 2019_08 - https://www.uniprot.org/).

### Synteny and comparative genome analysis

BLASTP (E-value 1×10^−5^) and MCScanX^83^ (-s 10) were used to identify homoeologous relationships between the two sub-genomes of fonio accession CM05836 and orthologous relationships between CM05836 and *Setaria italica*^*27*^, *Panicum miliaceum*^34^, *Panicum hallii*^*84*^, *Sorghum bicolor*^75^, *Zea mays*^76^, *Oryza sativa*^77^, *Brachypodium distachyon*^85^, *Hordeum vulgare*^86^, *Triticum aestivum*^30^ and *Aegilops tauschii*^87^. For all the comparisons, fonio sub-genomes A and B were analysed independently and only 1:1 and 1:2 relationships were retained according to the ploidy of the compared genomes.

To estimate the divergence time, we calculated the synonymous substitution rates (Ks) for each homoeologous and orthologous pair. Briefly, nucleotide sequences were aligned with clustalW^88^ and the Ks were calculated using the CODEML program in PAML^89^. Then, time of divergence events was calculated using T = Ks/2λ, based on the clock-(λ) estimated for grasses of 6.5 × 10^−9^ ^90^. To analyse genome dominance, the quantification of transcript abundance and expression were calculated using RSEM v1.3.1^91^ and genes were considered as expressed when expression was >0.5 TPM in at least one tissue. To study the homoeolog expression bias, we standardize the relative expression of the A and B gene pairs according to Ramirez-Gonzalez et al^31^.

### Mapping of re-sequencing data and variant calling

For quality control of each sample, raw sequence reads were analyzed with the fastqQC tool-v0.11.7 and low quality reads were filtered with trimmomatic-v0.38^71^ using the following criteria: LEADING:20; TRAILING:20; SLIDINGWINDOW:5:20 and MINLEN:50. The filtered paired-end reads were then aligned for each sample individually against the CM05836 reference assembly using BWA-MEM (v0.7.17-r1188)^53^ followed by sorting and indexing using samtools (v1.6)^92^. Alignment summary, base quality score and insert size metrics were collected and duplicated reads were marked and read groups were assigned using the Picard tools (http://broadinstitute.github.io/picard/). Variants were identified with GATK-v3.8^93^ using the – emitRefConfidence function of the HaplotypeCaller algorithm to call SNPs and InDels for each accession followed by a joint genotyping step performed by GenotypeGVCFs. To obtain high confidence variants, we excluded SNPs and InDels with the VariantFiltration function of GATK with the criteria: QD < 2.0; FS > 60.0; MQ < 40.0; MQRankSum < −12.5; ReadPosRankSum < - 8.0 and SOR > 4.0. The complete automated pipeline has been compiled and is available on github (https://github.com/IBEXCluster/IBEX-SNPcaller).

A total of 36.5 million variants were called. Raw variants were filtered using GATK-v3.8^93,94^ and VCFtools v0.1.17^95^. Variants located on chromosome unanchored, InDels, ‘SNP clusters’ defined as three or more SNPs located within 10 bp, missing data >10%, low and high average SNP depth (14 ≤ DP ≥ 42), and accessions having more than 33% of missing data were discarded. Only biallelic SNPs were retained to perform further analyses representing a final VCF file of 11,046,501 SNPs (Supplementary Table 12). These variants were annotated using snpEff v4.3^96^ with the CM05836 gene models.

### Genetic diversity and population structure

Genetic diversity and population structure analyses were performed using *D. exilis* and *D. longiflora* accessions together, or with *D. exilis* accessions alone. PCA and individual ancestry coefficients estimation were performed using the R package LEA^97^. The projections of the first PCA axis to geographic maps were performed following Cubry et al.^98^ and visualized using the Kriging package in R^99^. For ancestry coefficients analyses, the snmf function was used with ten independent runs for each *K* from *K* = 2 to *K* = 10. The optimal *K*, indicating the most likely number of ancestral populations, was determined with the cross-validation error rate. For the two analyses, we considered only SNPs present in at least two accessions. We also studied geographic distribution of private SNPs^98^ (i.e. SNP present only once in a single accession). Genome-wide pairwise linkage disequilibrium (LD) was estimated independently for *D. exilis* (2,617,322 SNPs) and *D. longiflora* (9,839,152 SNPs). Linkage disequilibrium (LD) decay (r^2^) was calculated using the PopLDdecay v.3.40 tool^100^ in a 500 kb distance and plotted using the ggplot2 package in R^66^.

### Association of climate, geography and social factors with the population genetics structure

Natural (i.e., climate, geography) and human (i.e., ethnic and linguistic) factors were tested for an association with the genetic structure of *D. exilis* accessions. Bioclimate data and ethnic and linguistic data were extracted for each of the 166 re-sequenced *D. exilis* accessions from WorldClim version 2^101^, passport data (ethnic groups), and Ethnologue version 16 (language, https://www.ethnologue.com/). The associations between the genetic structure and different factors were evaluated statistically using Pearson’s correlation coefficient test for the bioclimatic and geographic data with the R package devtools (https://devtools.r-lib.org/) (function cor). For the associations of social factors, two-sided Mantel tests (ecodist package - https://rdrr.io/cran/ecodist/) were performed between the genetic distance matrix of *D. exilis* and the dissimilarity matrix of social factors as inputs. Fisher’s exact tests (stat R package) were performed with the structure data at *K*=6 and accessions having ancestry thresholds of >70% as input. We used analysis of covariance (ANCOVA) to test for the impact of ethnolinguistic factors on the first PCA coordinates while controlling for other covariates: climatic and geographic variables.

### Demographic history reconstruction

A Sequentially Markovian Coalescent based approach^102,103^ was used to infer past evolution of the effective population size of *D. exilis* and *D. longiflora*. This analysis was made with the smc++^104^ software and additional customized scripts (https://github.com/Africrop/fonio_smcpp). We excluded SNPs that were not usable according to msmc-tools advices (https://github.com/stschiff/msmc-tools). In order to assess the impact of population structure on our results, additional analyses were done with all nine individuals from the genetically closest group of *D. longiflora* and also with the six groups of *D. exilis* based on the genetic structure. The smc++ vcf2smc tools was used to generate the input file and smc++ inference was performed for the different sets of genotypes using the smc++ cv command, considering 200 and 25,000 generations lower and upper time points boundaries respectively. All other options were set to default. We considered a generation time of one year and a mutation rate of 6.5 × 10^−9^. Different sets of distinguished lineages were considered for the different sets of individuals. For the random samples of *D. exilis* accessions, we randomly selected 20 lineages as distinguished ones, while for the other sets, all the samples were considered as distinguished. Graphical representation was made using the ggplot2^66^ package for R.

### Genome scanning for selection signals

Detection of selection was performed independently with three different methods: (1) Weir and Cockerham F_ST_; (2) nucleotide diversity ratio (π) were calculated between *D. exilis* and *D. longiflora* using VCFtools version 0.1.17^95^ with sliding windows of 50 kb every 10 kb; (3) SweeD version 3.3.1^37^ was used on *D. exilis* to detect signatures of selective sweeps based on the composite likelihood ratio (CLR) test within non-overlapping intervals of 3,000 bp along each chromosome. We grouped CLR peak positions into one common window if two or several peaks were less than 100 kb apart. Candidate genomic regions were selected for each method based on top 1% values. A list comprising 34 well characterized domestication genes (Supplementary data 3) from major crops was selected and compared by BLAST against gene models and the genome assembly of *D. exilis* accession CM05836. Each putative orthologous gene was validated by local alignment using ClustalW^105^. We considered the genes as orthologous when ≥ 70% of the coding sequence (CDS) length hit the genomic sequence of *D. exilis* with at least 70% identity. Validated orthologous genes were crossed with genomic regions under selection using bedtools v2.26.0^106^. To assess the effect of Δ*DeSh1* on seeds shattering, fonio panicles with three racemes were collected from mature plants and placed in 15ml tubes. Individual tubes were shaken using a Geno Grinder tissue homogenizer (SPEX SamplePrep, Metuchen, NJ) at 1,350 rpm for 20 seconds and the percentage of shattering was calculated. ANOVA test was performed with the MVApp (https://mvapp.kaust.edu.sa/)^107^.

### Data availability

The raw sequencing data used for *de novo* whole-genome assembly, the raw bionano map, the CM05836 genome assembly, the RNA-seq data for the annotation and the 183 re-sequenced accessions of *D. exilis* and *D. longiflora* for the population genomics analysis are available on EBI-ENA under the study number PRJEB36539.

The annotation of the CM05836 genome, the Tritex assembly and the VCF file are available on the DRYAD database under the doi:10.5061/dryad.2v6wwpzj0.

