## Supplementary Information for "Fonio millet genome unlocks African orphan crop diversity for agriculture in a changing climate"

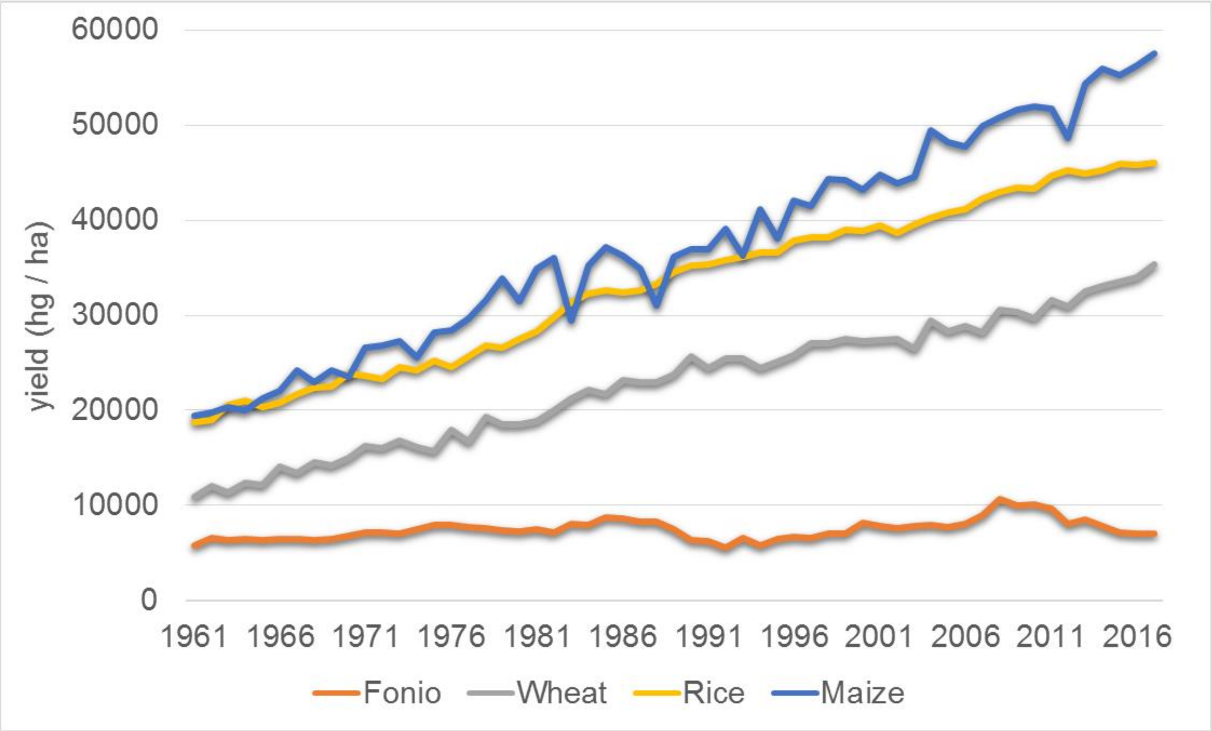


**Supplementary Fig. 1.** Yields (in hg / ha) of fonio, wheat, rice, and maize from 1961 to 2016. Data were collected from the Food and Agriculture Organization Corporate Statistical Database (FAOSTAT).

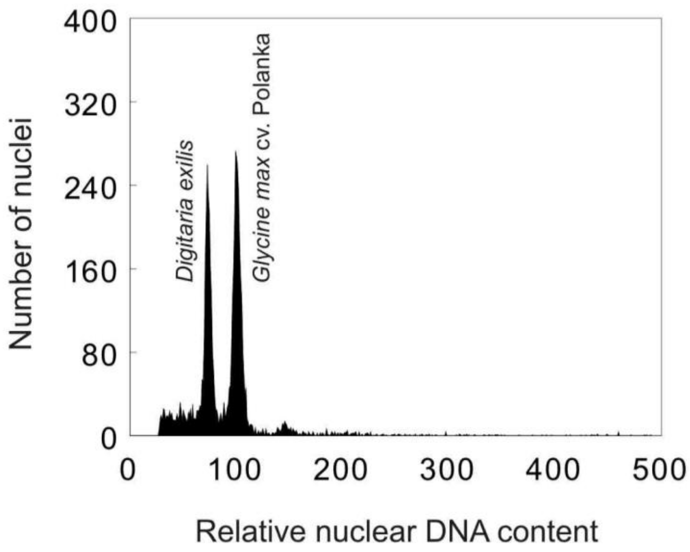


**Supplementary Fig. 2.** Estimation of nuclear genome size of *Digitaria exilis* using flow cytometry. The histogram of relative nuclear DNA content was obtained after flow cytometric analysis of propidium-iodide stained nuclei of *D. exilis* accession CM05836 and soybean (*Glycine max* cv. Polanka, 2C=2.5 pg), which served as the internal reference standard. 2C nuclear DNA content of *D. exilis* was estimated at 1.826 pg (± 0.004 s.d.).

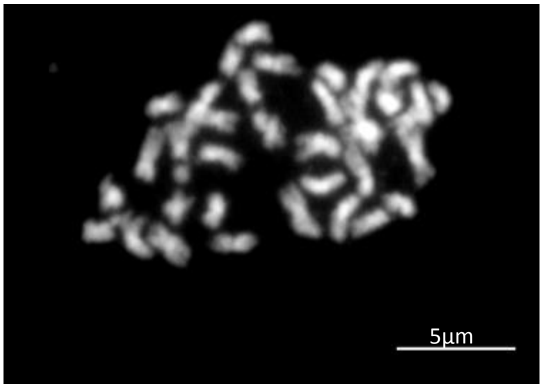


**Supplementary Fig. 3.** Mitotic metaphase plate of *D. exilis* accession CM05836 (2n = 4x = 36) showing size differences between the chromosomes. The chromosomes were stained by DAPI and the image is shown in white pseudocolor.

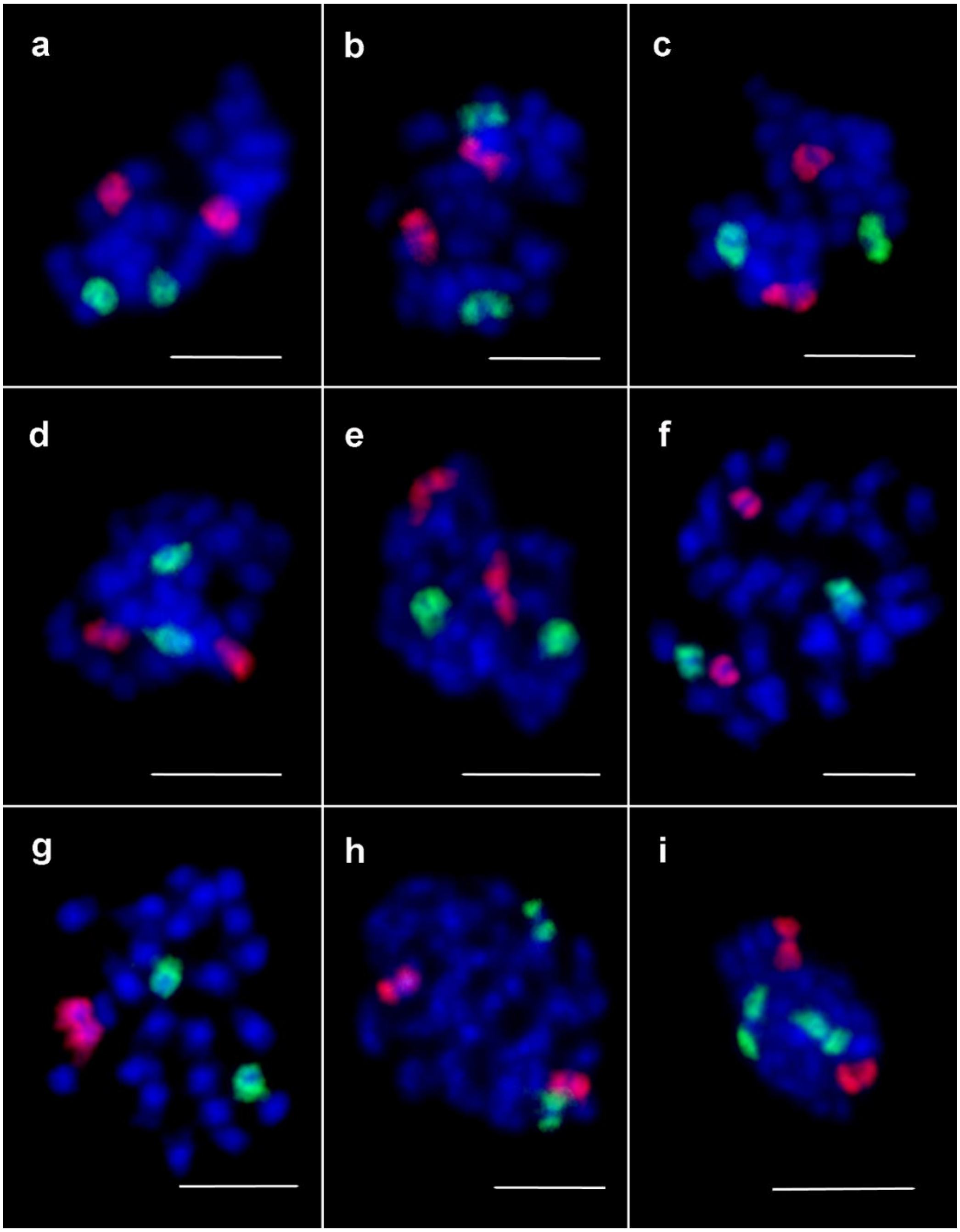


**Supplementary Fig. 4**. Oligo painting FISH on mitotic metaphase chromosomes of *Digitaria exilis* accession CM05836 (2n = 4x = 36). Each panel shows pairs of homoeologous chromosomes hybridized with 6-FAM (green) or CY3 (red) labeled painting probes. The oligomer probes were designed based on the assembled pseudomolecule sequences: **a,** pseudomolecules 1A (green) and 1B (red); **b,** pseudomolecules 2A (green) and 2B (red); **c,** pseudomolecules 3A (green) and 3B (red); **d,** pseudomolecules 4A (green) and 4B (red); **e,** pseudomolecules 5A (green) and 5B (red); **f,** pseudomolecules 6A (green) and 6B (red); **g,** pseudomolecules 7A (green) and 7B (red); **h,** pseudomolecules 8A (green) and 8B (red); and **i,** pseudomolecules 9A (green) and 9B (red). Chromosomes were counterstained with DAPI (blue). Scale bar = 5 µm.

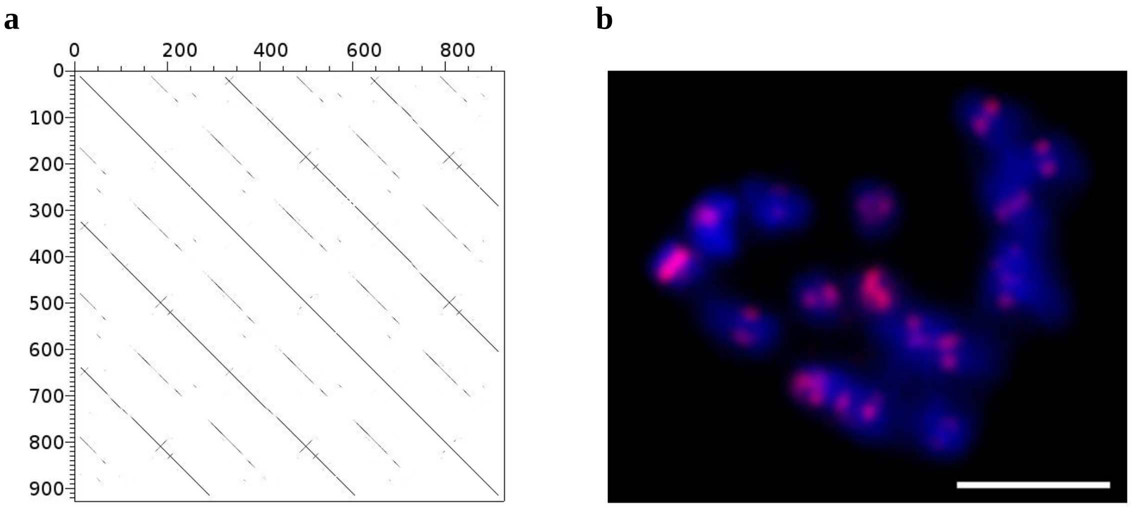


**Supplementary Fig. 5.** Identification of a centromeric repeat of *Digitaria exilis*. **a,** Dot plot analysis of tandem repeat CL10. **b,** FISH with a CY3 (red)- labeled probe for CL10 on mitotic metaphase plate revealed preferential localization to centromeric regions of all chromosomes. The chromosomes were counterstained with DAPI (blue). Scale bar = 5 µm.

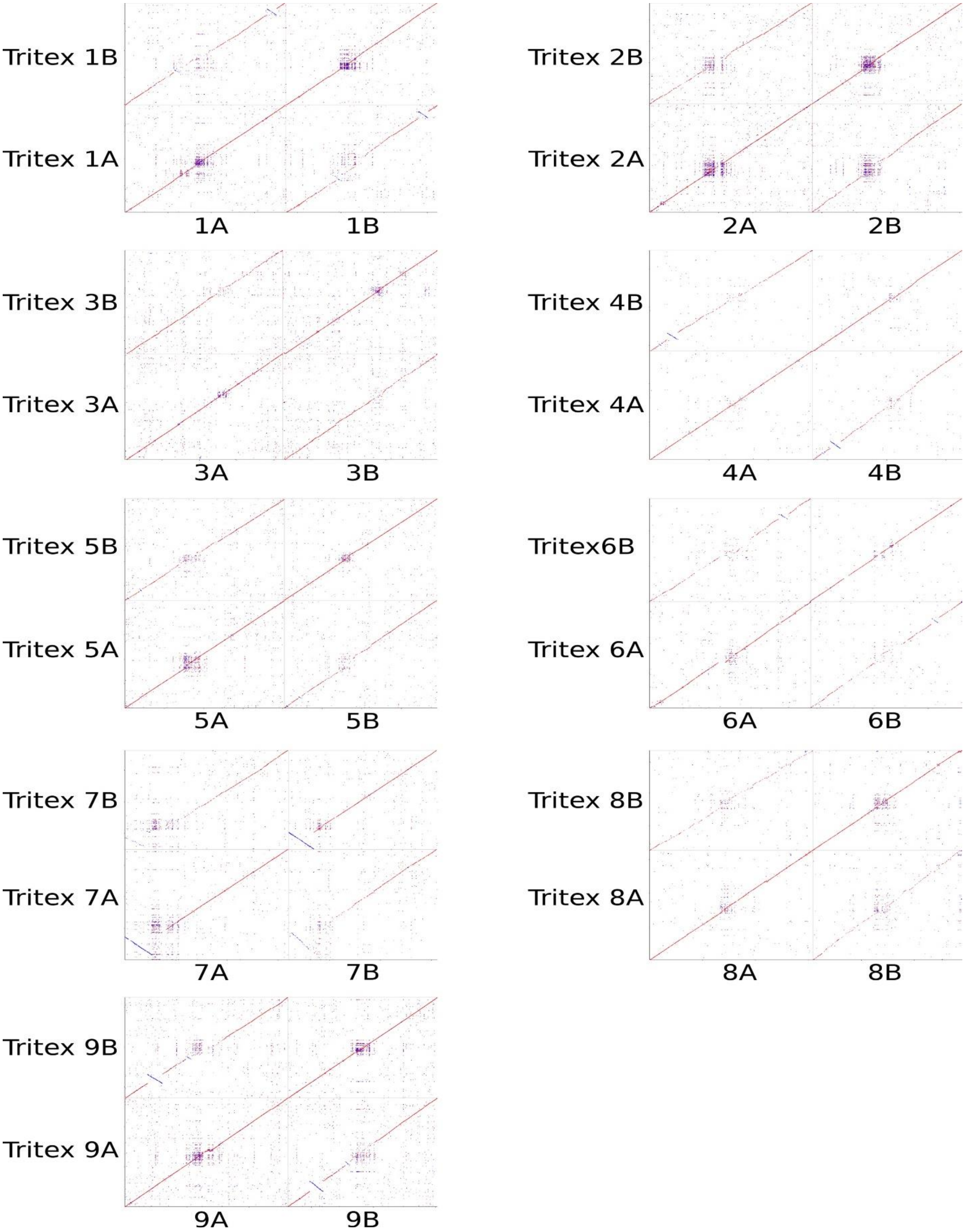


**Supplementary Fig. 6.** Comparison of *Digitaria exilis* CM05836 chromosomes assembled with DeNovoMAGIC3 and Tritex.

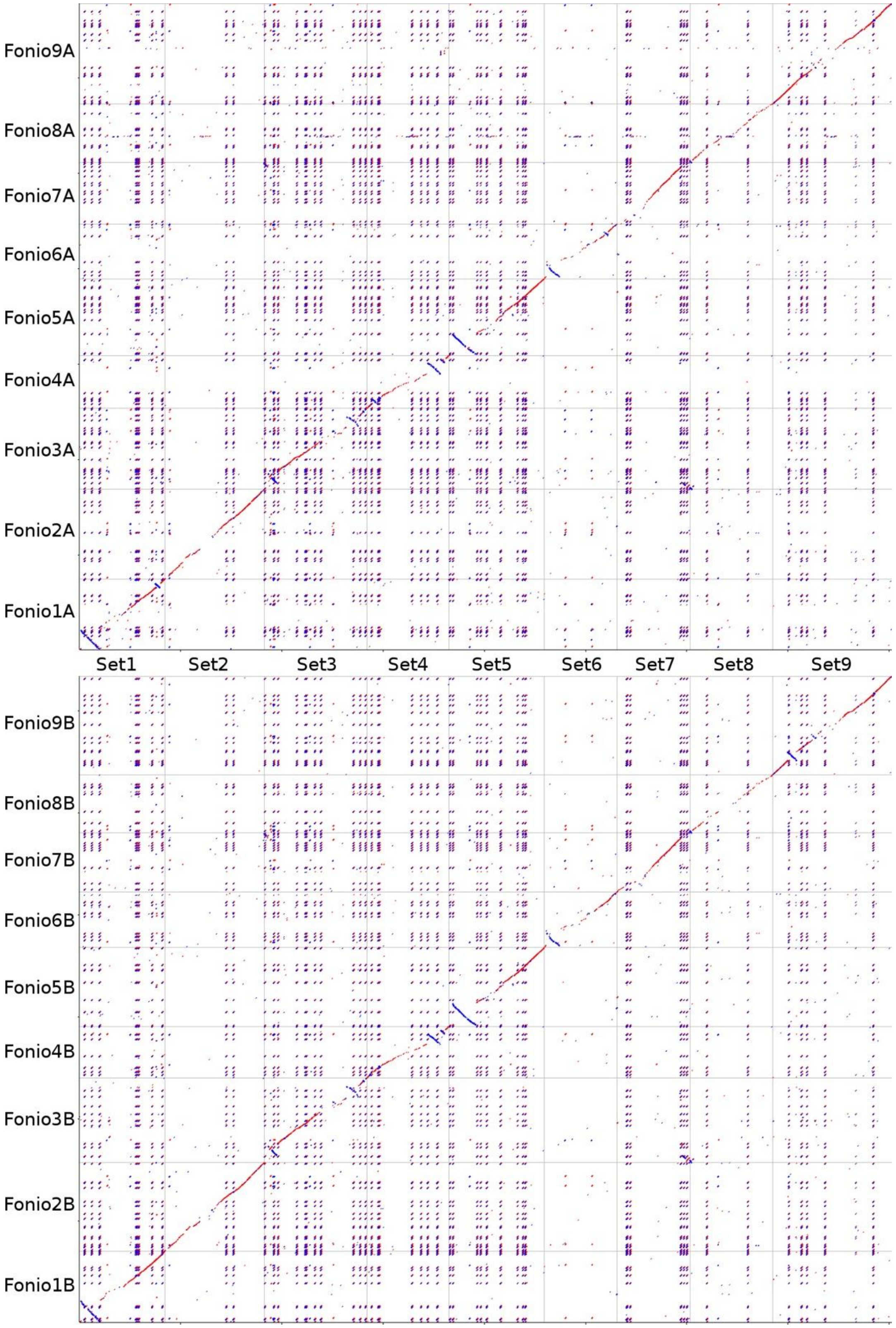


**Supplementary Fig. 7**. Comparison of *Digitaria exilis* CM05836 sub-genomes against *Setaria italica* (Set) shows syntenic relationship between the two species.

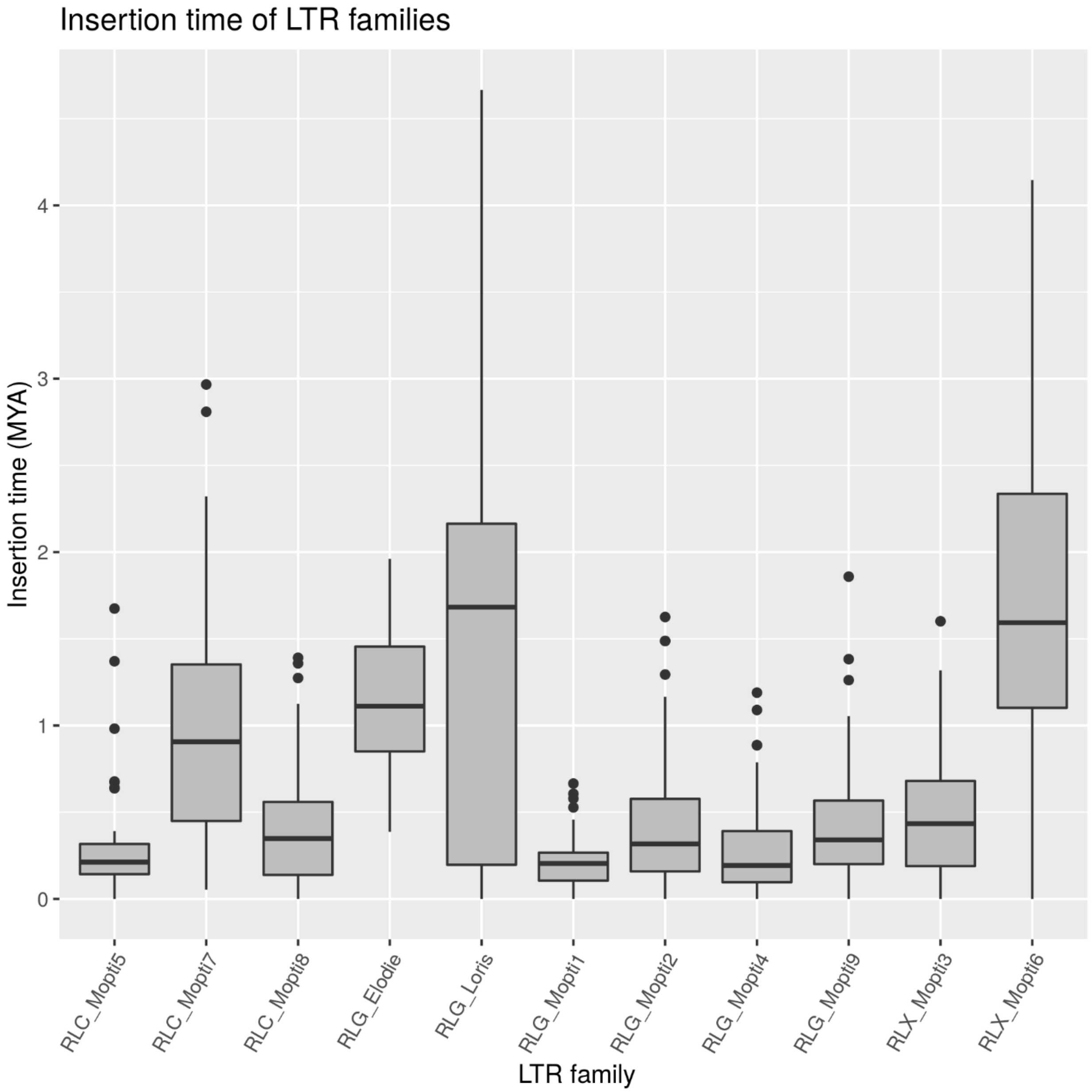


**Supplementary Fig. 8.** Insertion times of 11 full-length long terminal repeat retrotransposon (fl-LTR-RT) families in *Digitaria exilis* CM05836. Box plots of insertion time (million years ago MYA) for the 11 families with 30 intact copies are shown. The approximate insertion dates were calculated using the evolutionary distance between two LTR sequences of the same fl-LTR-RT. Clusters RLG_Loris and RLG_Elodie showed a differentiation between the two sub-genomes.

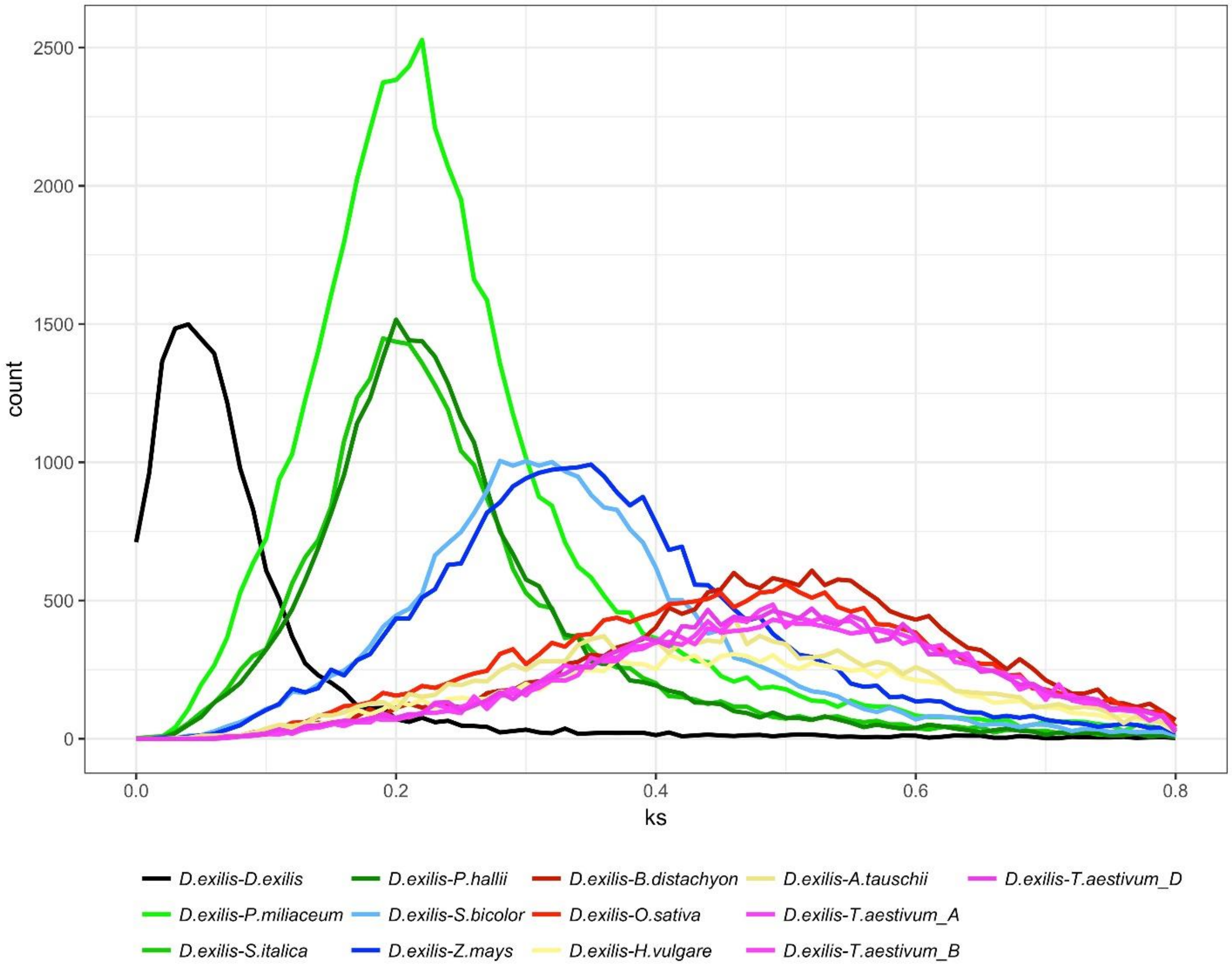


**Supplementary Fig. 9.** Ks distribution of homoeologous genes within *Digitaria exilis* and orthologous genes between *Digitaria exilis* and other monocots. Time of divergence events were calculated using the formula T = Ks/2λ, based on the clock-(λ) estimated for grasses of $6.5 \times 10^{-9}$.

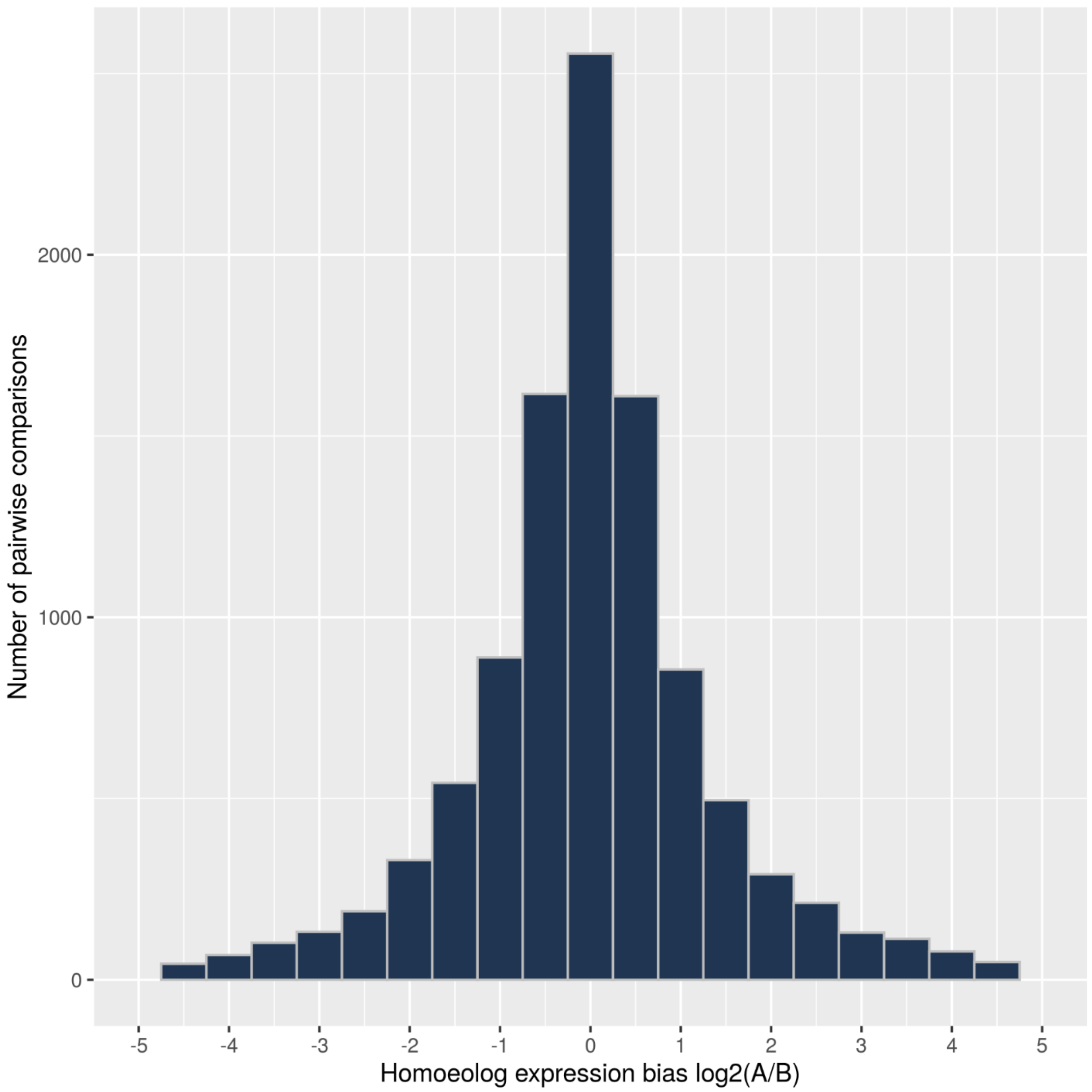


**Supplementary Fig. 10.** Homoeolog expression bias in *D. exilis* A and B sub-genomes considering all homoeologous gene pairs in flag-leaf, panicle, seedling and grain tissues. Values >0 indicate a bias toward the A sub-genome while values <0 indicate a bias toward the B sub-genome.

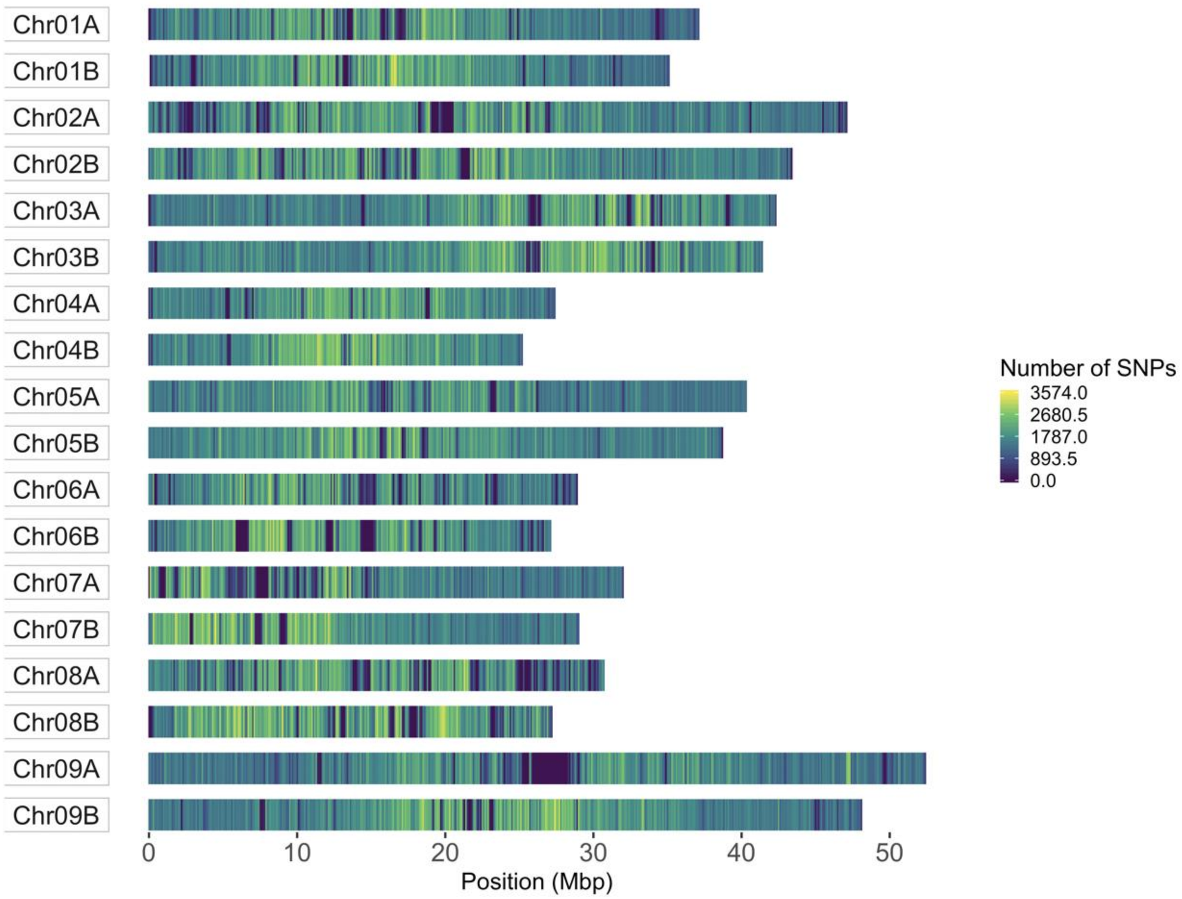


**Supplementary Fig. 11.** SNP density across the 18 *D. exilis* chromosomes calculated in a bin sizes of 100 kb.

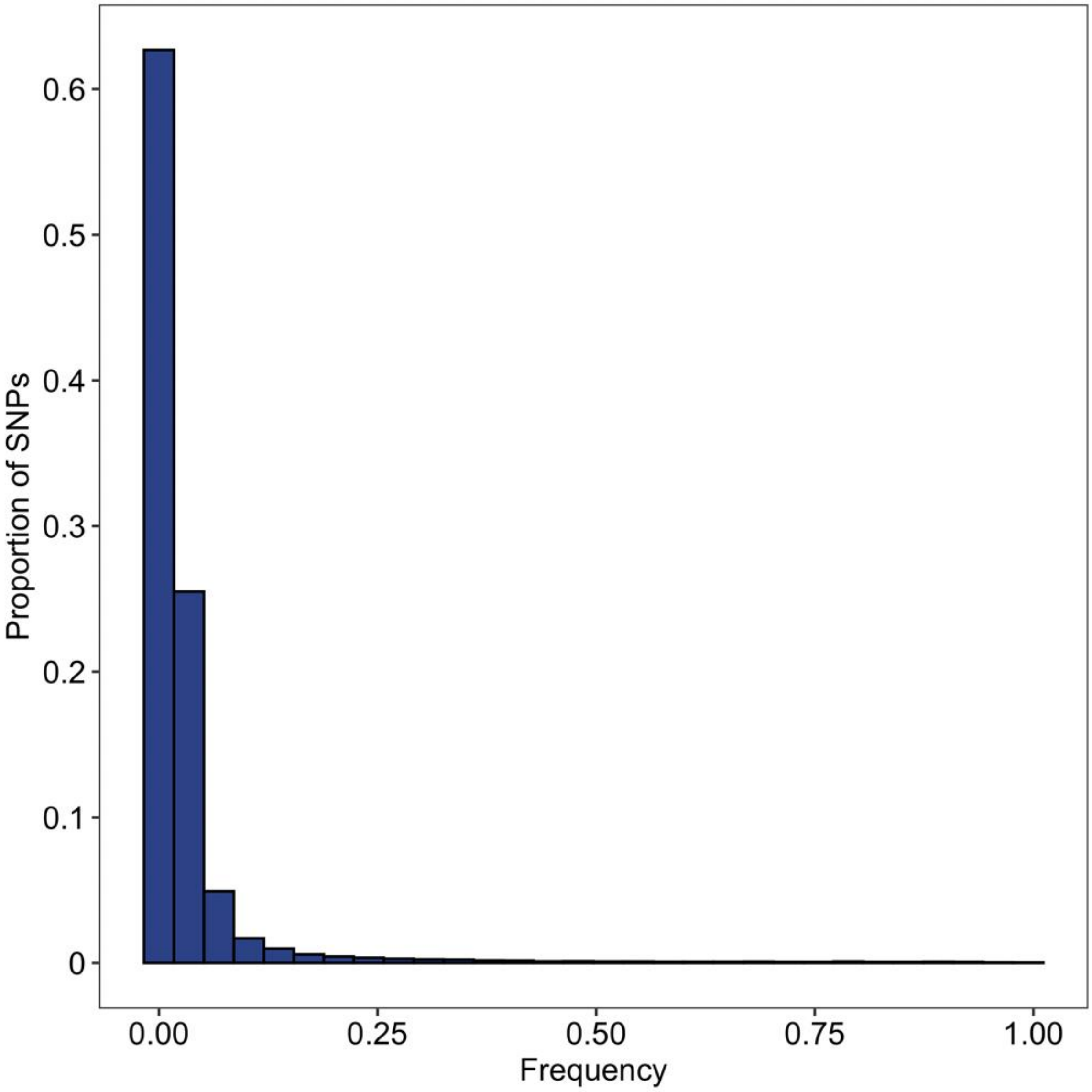


**Supplementary Fig. 12.** Site frequency spectrum of re-sequenced *D. exilis* and *D. longiflora* accessions reveals that most SNPs have a low minor allele frequency.

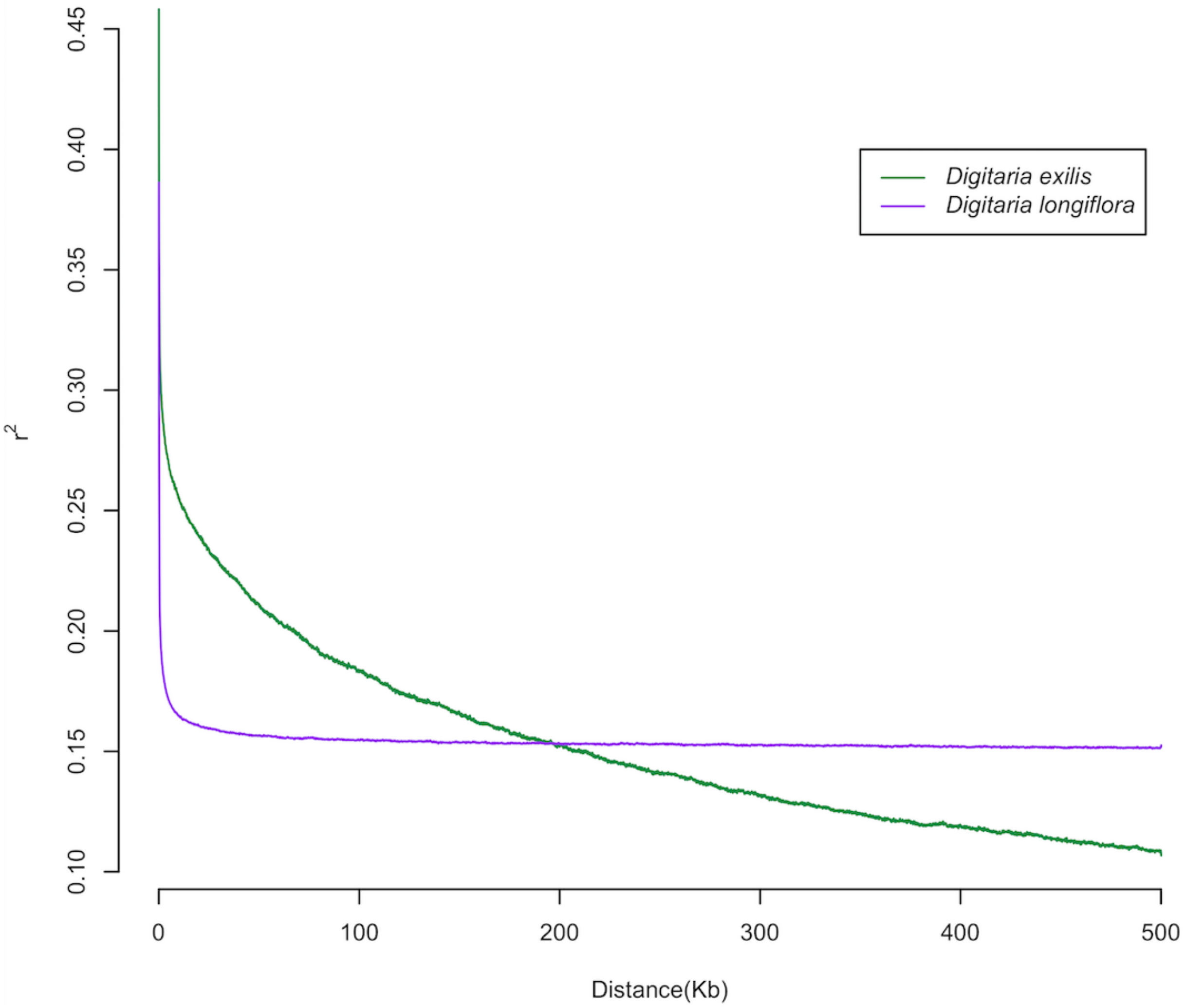


**Supplementary Fig. 13.** Genome-wide decay of linkage disequilibrium (LD).

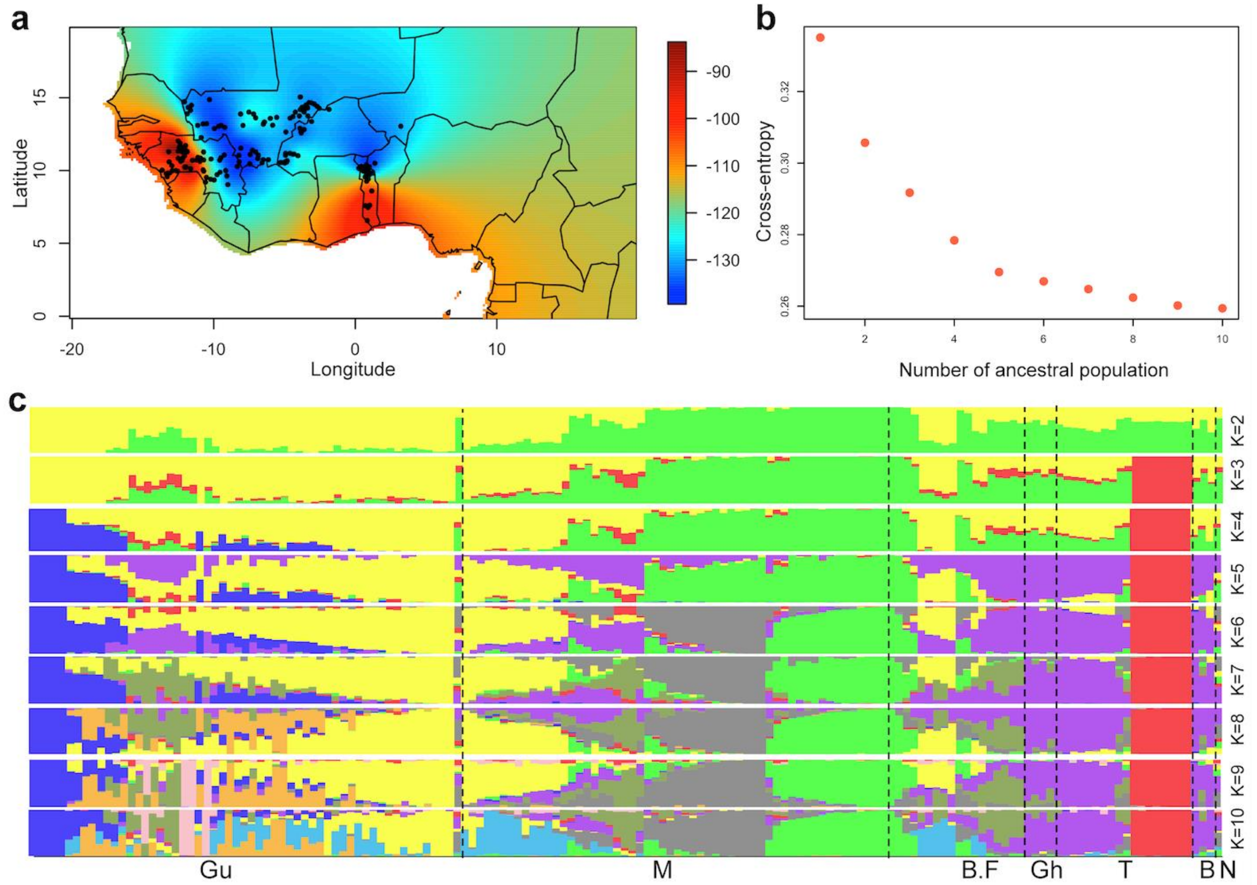


**Supplementary Fig. 14.** Population structure analyses of fonio. **a**, Geographical projection of the first PCA axis for *D. exilis* accessions. High values (red) indicate higher genetic relatedness of the respective *D. exilis* accessions to *D. longiflora*. Each accession is represented by a black dot. **b**, Cross-entropy for structure analysis of *D. exilis* samples. The cross entropy was calculated for a number of ancestral populations ranging from *K*=1 to *K*=10. **c**, Population structure (from *K*=2 to *K*=10) of *D. exilis* accessions. Each vertical bar represents an accession and the bars are filled by colors representing the likelihood of membership to each ancestry. The samples are ordered from west to east; Guinea (Gu), Mali (M), Burkina Faso (B.F), Ghana (Gh), Togo (T), Benin (B), and Niger (N).

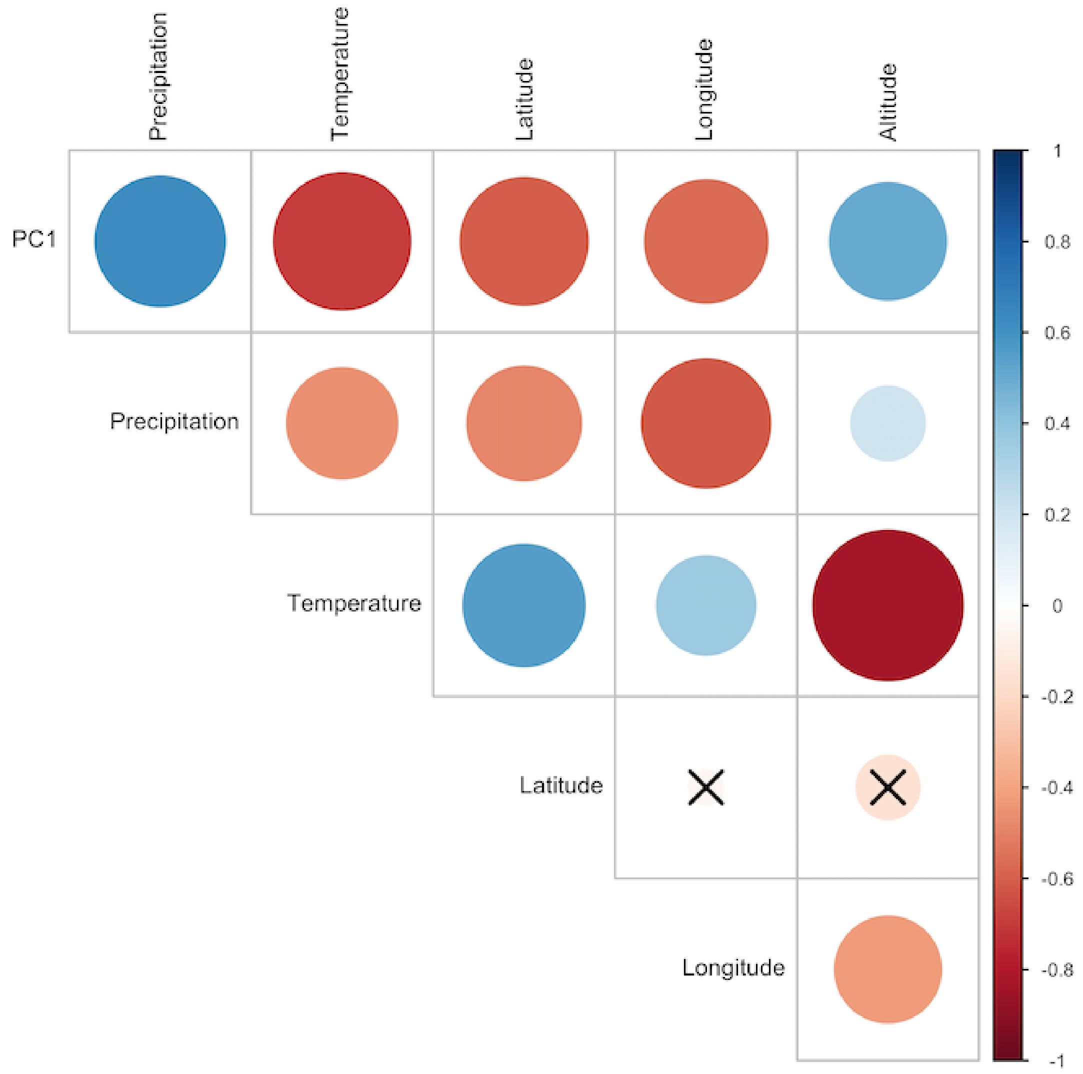


**Supplementary Fig. 15.** Pearson's correlation coefficients between the first principal coordinate (PC1) and climatic and geographic factors. The sizes of the circles reflect the strength of the correlation. Blue = positive correlation, red = negative correlation. The non-significant correlations, with a p-value above 0.05 are indicated with a cross in the individual cells.

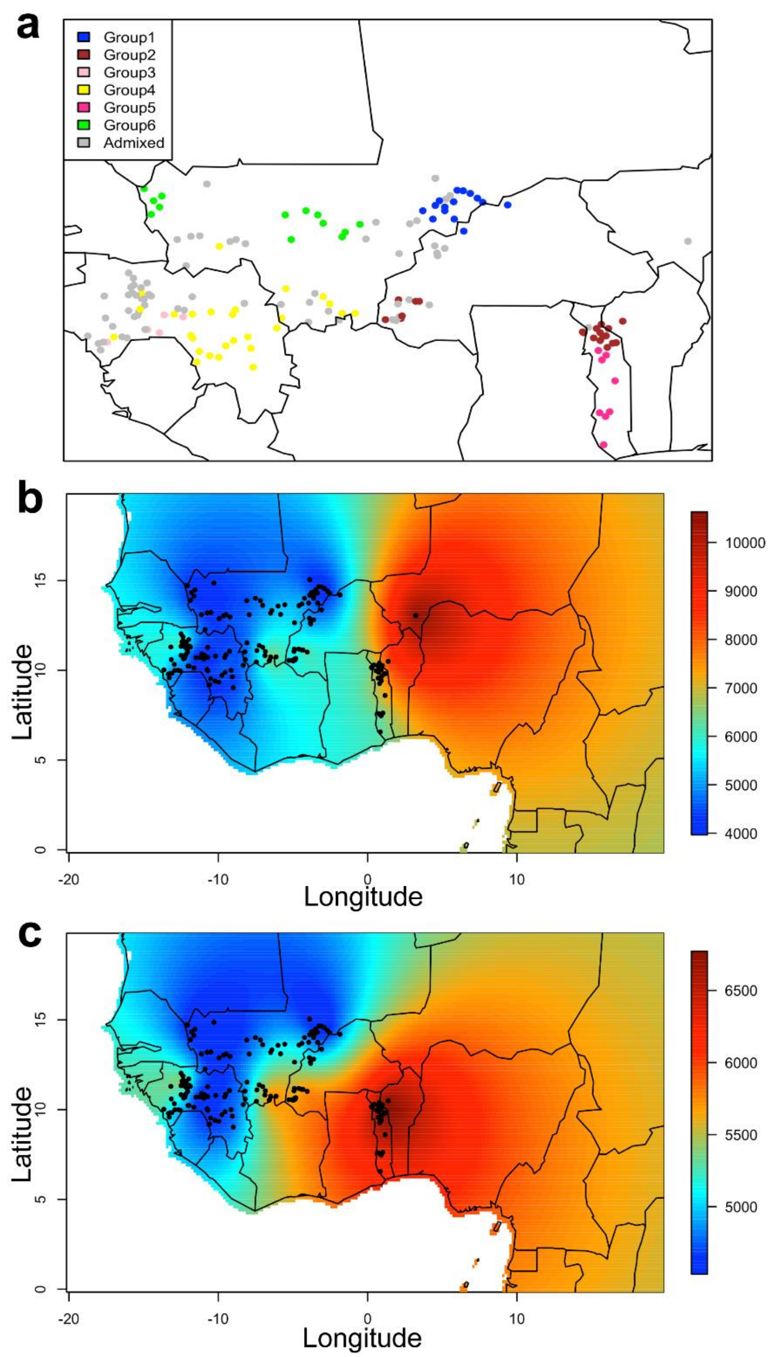


**Supplementary Fig. 16. a**, Map showing the geographic origin of *D. exilis* accessions colored based on the genetic structure at *K*=6. Gray dots represents the admixed accessions. **b**, and **c**, Geographic distribution of private SNPs (variants found only in one genotype) for all *D.exilis* accessions (**b**), and excluding one accession from Niger (**c**).

**Supplementary Table 1**. Raw sequencing data information

| Library type | Insert size | Approximate depth (coverage) | Total bases (Gb) | ENA sample number |
|---|---|---|---|---|
| PCR-free PE library (PE250x2) | 450-470 bp | 197x | 176.2 | ERS4280600 |
| PCR-free PE library (PE150x2) | 700- 800 bp | 124x | 110.5 | ERS4280601 |
| MP (Nextera™ MP Gel Plus) | 2-4 kb | 89x | 79.7 | ERS4280599 |
| MP (Nextera™ MP Gel Plus) | 5-7 kb | 80x | 71.6 | ERS4280602 |
| MP (Nextera™ MP Gel Plus) | 8-10 kb | 72x | 63.9 | ERS4280603 |
| 10x Genomics™ Chromium™ | N/A | 84x | 74.6 | - |
| Hi-C | - | 122x | 109 | ERS4280610 – ERS4280605 |
| Bionano | - | 235x | 210.6 | ERZ1300513 |
| RNAseq seedling | | - | 32.2 | ERS4280607 |
| RNAseq grain | | - | 30.2 | ERS4280606 |
| RNAseq flag leaf | | - | 24.4 | ERS4280609 |
| RNAseq panicle | | - | 24.7 | ERS4280608 |

**Supplementary Table 2**. Statistics of the CM05836 assembly after Hi-C integration.

| | |
|---|---|
| Total length of assembly (Mb) | **701.680** |
| Number of super-scaffolds | **8,293** |
| Longest super-scaffold (Mb) | **49.06** |
| N50 (Mb) | **35.296** |
| N90 (Mb) | **25.563** |
| Number of gaps | **18,529** |
| Number of breaks made to input assembly by HiRise | **5** |
| Number of joints made by HiRise | **174** |

**Supplementary Table 3**. Statistics of the Bionano assembly.

| | Bionano Assembly |
|---|---|
| **Molecules** | |
| Molecules aligned to NRGene assembly | **739,153** |
| Filtered molecule N50 (>=150 kb) | **183** |
| **Assembly** | |
| Length of Bionano assembly (Mb) | **714.793** |
| Number of scaffolds | **8,406** |
| Longest scaffold (Mb) | **40.867** |
| N50 (Mb) | **21.104** |
| N90 (Mb) | **6.519** |
| Length of hybrid scaffold assembly(Mb) | **657.324** |
| Number of hybrid scaffolds | **39** |
| Longest hybrid scaffold (Mb) | **40.867** |
| N50 (Mb) | **22.646** |
| N90 (Mb) | **11.849** |
| **Conflict** | |
| Deletions | **412** |
| Insertions | **1,858** |
| Duplications | **5** |

**Supplementary Table 4**. Number of annotated transcripts per tissue and sub-genome.

| | sub-genome A | sub-genome B | unanchored |
|---|---|---|---|
| **flag leaf** | 15,073 | 14,998 | 1,191 |
| **grain** | 13,454 | 13,447 | 1,055 |
| **seedling** | 18,115 | 18,064 | 1,425 |
| **panicle** | 19,390 | 19,430 | 1,513 |

**Supplementary Table 5**. Number of transcripts expressed in multiple CM05836 tissues.

| | **sub-genome A** | **Sub-genomeB** | **unanchored** | **Total** | |
|---|---|---|---|---|---|
| **Never expressed** | 7,306 | 7,046 | 1,089 | 15,441 | **25.74%** |
| **Expressed in 1 tissue** | 3,231 | 3,190 | 312 | 6,733 | **74.26%** |
| **Expressed in 2 tissues** | 2,879 | 2,875 | 252 | 6,006 | |
| **Expressed in 3 tissues** | 4,225 | 4,237 | 340 | 8,802 | |
| **Expressed in 4 tissues** | 11,092 | 11,072 | 837 | 23,001 | |

**Supplementary Table 6**. Number of raw reads, total base pairs (Gb) and sequence coverage of *D. exilis* and *D. longiflora*. Coverage was calculated based on the CM05836 assembly size. Samples in blue were excluded from analyses because of missing data.

| Accession | Nr. of raw read pairs | Total base pairs (Gb) | Coverage |
|---|---|---|---|
| CM03380 | 129,530,462 | 38.9 | 54.3 |
| CM03382 | 121,455,096 | 36.4 | 50.8 |
| CM03390 | 107,058,717 | 32.1 | 44.8 |
| CM03394 | 122,970,585 | 36.9 | 51.5 |
| CM03396 | 97,519,029 | 29.3 | 40.9 |
| CM03400 | 113,926,105 | 34.2 | 47.7 |
| CM03403 | 117,281,810 | 35.2 | 49.1 |
| CM03406 | 89,702,848 | 26.9 | 37.5 |
| CM03409 | 107,688,516 | 32.3 | 45.1 |
| CM03418 | 111,133,919 | 33.3 | 46.5 |
| CM03423 | 124,051,104 | 37.2 | 51.9 |
| CM03424 | 102,702,604 | 30.8 | 43.0 |
| CM03430 | 129,312,103 | 38.8 | 54.2 |
| CM03431 | 101,053,499 | 30.3 | 42.3 |
| CM03434 | 110,830,696 | 33.2 | 46.3 |
| CM03437 | 114,075,345 | 34.2 | 47.7 |
| CM03438 | 113,327,472 | 34.0 | 47.5 |
| CM03439 | 105,740,268 | 31.7 | 44.2 |
| CM04487 | 108,642,286 | 32.6 | 45.5 |
| CM04489 | 112,834,857 | 33.9 | 47.3 |
| CM04493 | 111,592,219 | 33.5 | 46.8 |
| CM04495 | 106,019,039 | 31.8 | 44.4 |
| CM05734 | 95,007,889 | 28.5 | 39.8 |
| CM05736 | 99,371,418 | 29.8 | 41.6 |
| CM05737 | 108,145,394 | 32.4 | 45.2 |
| CM05740 | 114,662,297 | 34.7 | 48.4 |
| CM05741 | 125,628,296 | 37.7 | 52.6 |
| CM05742 | 116,207,696 | 35.1 | 49.0 |
| CM05743 | 102,129,790 | 30.6 | 42.7 |
| CM05744 | 114,417,387 | 34.3 | 47.9 |
| CM05746 | 100,579,569 | 30.2 | 42.2 |
| CM05750 | 112,770,059 | 33.8 | 47.2 |
| CM05754 | 101,017,430 | 30.3 | 42.3 |
| CM05757 | 107,847,260 | 32.4 | 45.2 |
| CM05760 | 146,208,305 | 43.9 | 61.3 |

| | | | |
|---|---|---|---|
| CM05761 | 116,158,639 | 34.8 | 48.6 |
| CM05767 | 99,117,778 | 29.7 | 41.5 |
| CM05770 | 87,097,747 | 26.1 | 36.4 |
| CM05771 | 90,158,763 | 27.0 | 37.7 |
| CM05772 | 96,955,246 | 29.1 | 40.6 |
| CM05775 | 92,385,118 | 27.7 | 38.7 |
| CM05780 | 96,829,037 | 29.0 | 40.5 |
| CM05792 | 97,533,677 | 29.3 | 40.9 |
| CM05793 | 109,973,076 | 33.0 | 46.1 |
| CM05796 | 99,606,969 | 29.9 | 41.7 |
| CM05798 | 108,330,985 | 32.5 | 45.4 |
| CM05802 | 89,133,336 | 26.7 | 37.3 |
| CM05803 | 88,659,420 | 26.6 | 37.1 |
| CM05806 | 105,285,367 | 31.6 | 44.1 |
| CM05808 | 116,007,622 | 34.8 | 48.6 |
| CM05810 | 99,685,620 | 29.9 | 41.7 |
| CM05811 | 108,230,928 | 32.5 | 45.4 |
| CM05815 | 97,840,195 | 29.4 | 41.0 |
| CM05819 | 104,895,469 | 31.5 | 44.0 |
| CM05824 | 116,103,893 | 34.8 | 48.6 |
| CM05827 | 106,074,007 | 31.8 | 44.4 |
| CM05830 | 114,691,379 | 34.4 | 48.0 |
| CM05834 | 105,237,809 | 31.6 | 44.1 |
| CM05835 | 94,427,566 | 28.3 | 39.5 |
| CM05836 | 121,132,900 | 36.6 | 51.1 |
| CM05836-D | 111,402,466 | 33.7 | 47.0 |
| CM05839 | 107,877,804 | 32.4 | 45.2 |
| CM05840 | 109,891,555 | 33.0 | 46.1 |
| CM05841 | 92,687,779 | 27.8 | 38.8 |
| CM05842 | 99,316,439 | 29.8 | 41.6 |
| CM05843 | 118,584,430 | 35.6 | 49.7 |
| CM05844 | 109,003,223 | 32.7 | 45.6 |
| CM05847 | 96,949,493 | 29.1 | 40.6 |
| CM05849 | 97,551,329 | 29.3 | 40.9 |
| CM05853 | 103,537,228 | 31.1 | 43.4 |
| CM05854 | 101,827,075 | 30.5 | 42.6 |
| CM05855 | 103,594,323 | 31.1 | 43.4 |
| CM05856 | 113,883,533 | 34.5 | 48.1 |
| CM05857 | 104,876,036 | 31.5 | 44.0 |
| CM05858 | 114,451,268 | 34.3 | 47.9 |
| CM05863 | 105,904,412 | 31.8 | 44.4 |

| | | | |
|---|---|---|---|
| CM05865 | 100,686,174 | 30.2 | 42.2 |
| CM05869 | 116,770,456 | 35.0 | 48.9 |
| CM05870 | 117,403,534 | 35.2 | 49.1 |
| CM06496 | 108,023,288 | 32.7 | 45.6 |
| CM06501 | 107,957,703 | 32.4 | 45.2 |
| CM06505 | 110,254,970 | 33.1 | 46.2 |
| CM06510 | 98,810,685 | 29.6 | 41.3 |
| CM06511 | 105,976,725 | 31.8 | 44.4 |
| CM06513 | 105,697,489 | 31.7 | 44.2 |
| CM07226 | 114,283,176 | 34.3 | 47.9 |
| CM07234 | 113,061,844 | 33.9 | 47.3 |
| CM07244 | 113,259,206 | 34.0 | 47.5 |
| CM07246 | 107,502,998 | 32.3 | 45.1 |
| CM07249 | 137,973,150 | 41.4 | 57.8 |
| CM07258 | 117,580,556 | 35.3 | 49.3 |
| CM07261 | 105,040,951 | 31.5 | 44.0 |
| CM07262 | 130,515,966 | 39.2 | 54.7 |
| CM07264 | 124,523,838 | 37.4 | 52.2 |
| CM07266 | 94,175,090 | 28.3 | 39.5 |
| CM07268 | 114,829,434 | 34.4 | 48.0 |
| CM07269 | 95,345,749 | 28.6 | 39.9 |
| CM07270 | 128,603,417 | 38.6 | 53.9 |
| CM07272 | 116,616,988 | 35.0 | 48.9 |
| CM07277 | 111,666,263 | 33.5 | 46.8 |
| CM07280 | 92,760,311 | 27.8 | 38.8 |
| CM07281 | 125,452,545 | 37.6 | 52.5 |
| CM07283 | 101,607,547 | 30.5 | 42.6 |
| CM07285 | 90,868,945 | 27.3 | 38.1 |
| CM07289 | 91,094,917 | 27.3 | 38.1 |
| CM07292 | 87,091,276 | 26.1 | 36.4 |
| CM07295 | 100,698,705 | 30.2 | 42.2 |
| CM07300 | 94,991,516 | 28.5 | 39.8 |
| CM07310 | 95,459,056 | 28.6 | 39.9 |
| CM07317 | 91,134,013 | 27.3 | 38.1 |
| CM07319 | 90,269,759 | 27.1 | 37.8 |
| CM07321 | 99,501,199 | 29.9 | 41.7 |
| CM07327 | 96,986,970 | 29.1 | 40.6 |
| CM07329 | 104,986,252 | 31.5 | 44.0 |
| CM07338 | 93,318,292 | 28.0 | 39.1 |
| CM07340 | 114,809,162 | 34.4 | 48.0 |
| CM07342 | 96,685,623 | 29.0 | 40.5 |

| | | | |
|---|---|---|---|
| CM07346 | 117,181,077 | 35.2 | 49.1 |
| CM07350 | 106,385,085 | 31.9 | 44.5 |
| CM07354 | 102,269,101 | 30.7 | 42.8 |
| CM07360 | 122,186,789 | 36.7 | 51.2 |
| CM07885 | 88,674,361 | 26.6 | 37.1 |
| CM07888 | 141,602,386 | 42.5 | 59.3 |
| CM07890 | 102,755,504 | 30.8 | 43.0 |
| CM07891 | 90,176,543 | 27.1 | 37.8 |
| CM07892 | 115,798,450 | 35.0 | 48.8 |
| CM07895 | 92,770,978 | 27.8 | 38.8 |
| CM07900 | 89,604,850 | 26.9 | 37.5 |
| CM07901 | 105,486,161 | 31.6 | 44.1 |
| CM07902 | 109,253,262 | 32.8 | 45.8 |
| CM07903 | 88,205,352 | 26.5 | 37.0 |
| CM07908 | 92,613,952 | 27.8 | 38.8 |
| CM07909 | 116,736,515 | 35.0 | 48.9 |
| CM08512 | 96,797,035 | 29.0 | 40.5 |
| CM08517 | 90,087,089 | 27.0 | 37.7 |
| CM08524 | 90,416,496 | 27.1 | 37.8 |
| CM08525 | 97,118,027 | 29.1 | 40.6 |
| CM08526 | 90,101,625 | 27.0 | 37.7 |
| CM08537 | 99,910,677 | 30.0 | 41.9 |
| CM08551 | 88,274,090 | 26.5 | 37.0 |
| CM08562 | 101,054,449 | 30.3 | 42.3 |
| CM08563 | 100,980,626 | 30.3 | 42.3 |
| CM08573 | 100,489,420 | 30.1 | 42.0 |
| CM08593 | 115,208,380 | 34.6 | 48.3 |
| CM08598 | 94,852,814 | 28.5 | 39.8 |
| CM08613 | 121,007,623 | 36.3 | 50.7 |
| CM08617 | 114,845,352 | 34.5 | 48.2 |
| CM08627 | 93,684,295 | 28.1 | 39.2 |
| CM08628 | 113,141,265 | 33.9 | 47.3 |
| CM08635 | 96,348,482 | 28.9 | 40.3 |
| CM08642 | 106,268,044 | 31.9 | 44.5 |
| CM08652 | 133,102,982 | 39.9 | 55.7 |
| CM08653 | 106,672,009 | 32.0 | 44.7 |
| CM08655 | 108,948,318 | 32.7 | 45.6 |
| CM08665 | 105,674,069 | 31.7 | 44.2 |
| CM08669 | 98,067,489 | 29.4 | 41.0 |
| CM08671 | 103,282,194 | 31.0 | 43.3 |
| CM08672 | 118,737,111 | 35.6 | 49.7 |

| | | | |
|---|---|---|---|
| CM08677 | 119,278,939 | 35.8 | 50.0 |
| CM08681 | 143,247,100 | 43.0 | 60.0 |
| CM08684 | 98,467,427 | 29.5 | 41.2 |
| CM12246 | 119,691,265 | 36.2 | 50.5 |
| CH1 | 112,506,107 | 34.0 | 47.5 |
| CH2 | 116,224,409 | 35.1 | 49.1 |
| CH3 | 124,088,536 | 37.5 | 52.4 |
| CH4 | 123,164,702 | 37.2 | 52.0 |
| 4349 | 37,791,196 | 10.0 | 14.0 |
| 67996 | 67,523,402 | 19.8 | 27.6 |
| If-Dlon1 | 59,937,518 | 17.3 | 24.2 |
| If-Dlon12 | 50,988,995 | 7.2 | 10.0 |
| L121 | 61,386,054 | 17.8 | 24.9 |
| L17 | 66,614,904 | 19.1 | 26.6 |
| L3027 | 57,475,984 | 16.8 | 23.4 |
| L4148 | 39,283,204 | 11.4 | 15.9 |
| L8249 | 36,137,687 | 10.6 | 14.7 |
| L9512 | 26,666,396 | 7.8 | 10.9 |
| L953 | 53,517,704 | 15.6 | 21.8 |
| M1070 | 48,565,006 | 12.9 | 18.0 |
| M861 | 54,196,416 | 15.7 | 21.9 |
| M865 | 45,462,023 | 13.2 | 18.4 |
| M883 | 58,308,861 | 16.9 | 23.6 |
| M905 | 51,810,076 | 15.0 | 20.9 |
| M949 | 67,901,376 | 19.8 | 27.7 |

**Supplementary Table 7**. Mapping rate of raw reads of each re-sequenced *D. exilis* and *D. longiflora* accession to the CM05836 reference assembly. Samples in blue were excluded from analyses because of missing data.

| Accession | Total number of raw reads | Number of mapped reads | Mapping rate |
|---|---|---|---|
| CM03380 | 259,060,924 | 228,203,469 | 88.09 |
| CM03382 | 242,910,192 | 214,725,124 | 88.40 |
| CM03390 | 214,117,434 | 188,807,070 | 88.18 |
| CM03394 | 245,941,170 | 217,215,887 | 88.32 |
| CM03396 | 195,038,058 | 171,976,760 | 88.18 |
| CM03400 | 227,852,210 | 201,428,175 | 88.40 |
| CM03403 | 234,563,620 | 207,236,020 | 88.35 |
| CM03406 | 179,405,696 | 158,235,344 | 88.20 |
| CM03409 | 215,377,032 | 189,140,329 | 87.82 |
| CM03418 | 222,267,838 | 196,002,971 | 88.18 |
| CM03423 | 248,102,208 | 218,477,110 | 88.06 |
| CM03424 | 205,405,208 | 181,307,384 | 88.27 |
| CM03430 | 258,624,206 | 228,271,052 | 88.26 |
| CM03431 | 202,106,998 | 178,325,138 | 88.23 |
| CM03434 | 221,661,392 | 195,406,290 | 88.16 |
| CM03437 | 228,150,690 | 201,158,715 | 88.17 |
| CM03438 | 226,654,944 | 198,823,904 | 87.72 |
| CM03439 | 211,480,536 | 186,391,555 | 88.14 |
| CM04487 | 217,284,572 | 191,699,500 | 88.23 |
| CM04489 | 225,669,714 | 199,323,170 | 88.33 |
| CM04493 | 223,184,438 | 197,099,165 | 88.31 |
| **CM04495** | 212,038,078 | 97,251,294 | 45.87 |
| CM05734 | 190,015,778 | 167,478,198 | 88.14 |
| CM05736 | 198,742,836 | 175,522,986 | 88.32 |
| CM05737 | 216,290,788 | 190,495,300 | 88.07 |
| CM05740 | 229,324,594 | 203,409,801 | 88.70 |
| CM05741 | 251,256,592 | 221,739,502 | 88.25 |
| CM05742 | 232,415,392 | 206,358,122 | 88.79 |
| CM05743 | 204,259,580 | 180,512,515 | 88.37 |
| CM05744 | 228,834,774 | 202,005,656 | 88.28 |
| CM05746 | 201,159,138 | 176,996,117 | 87.99 |
| CM05750 | 225,540,118 | 198,231,435 | 87.89 |
| CM05754 | 202,034,860 | 176,690,294 | 87.46 |
| **CM05757** | 215,694,520 | 85,458,624 | 39.62 |
| CM05760 | 292,416,610 | 255,475,124 | 87.37 |
| CM05761 | 232,317,278 | 203,968,882 | 87.80 |

| | | | |
|---|---|---|---|
| CM05767 | 198,235,556 | 174,455,815 | 88.00 |
| CM05770 | 174,195,494 | 153,768,478 | 88.27 |
| CM05771 | 180,317,526 | 158,531,060 | 87.92 |
| CM05772 | 193,910,492 | 168,653,894 | 86.98 |
| CM05775 | 184,770,236 | 162,553,908 | 87.98 |
| CM05780 | 193,658,074 | 170,062,427 | 87.82 |
| CM05792 | 195,067,354 | 172,331,664 | 88.34 |
| CM05793 | 219,946,152 | 194,445,565 | 88.41 |
| CM05796 | 199,213,938 | 175,559,736 | 88.13 |
| CM05798 | 216,661,970 | 192,121,545 | 88.67 |
| CM05802 | 178,266,672 | 157,077,197 | 88.11 |
| CM05803 | 177,318,840 | 156,697,151 | 88.37 |
| CM05806 | 210,570,734 | 185,976,667 | 88.32 |
| CM05808 | 232,015,244 | 202,335,334 | 87.21 |
| CM05810 | 199,371,240 | 176,078,110 | 88.32 |
| CM05811 | 216,461,856 | 191,404,684 | 88.42 |
| CM05815 | 195,680,390 | 172,783,063 | 88.30 |
| CM05819 | 209,790,938 | 186,043,694 | 88.68 |
| CM05824 | 232,207,786 | 205,093,066 | 88.32 |
| CM05827 | 212,148,014 | 187,724,146 | 88.49 |
| CM05830 | 229,382,758 | 202,989,950 | 88.49 |
| CM05834 | 210,475,618 | 185,917,010 | 88.33 |
| CM05835 | 188,855,132 | 167,687,418 | 88.79 |
| CM05836 | 242,265,800 | 216,481,300 | 89.36 |
| CM05836-D | 222,804,932 | 196,969,293 | 88.40 |
| CM05839 | 215,755,608 | 190,912,997 | 88.49 |
| CM05840 | 219,783,110 | 195,074,162 | 88.76 |
| CM05841 | 185,375,558 | 164,492,091 | 88.73 |
| CM05842 | 198,632,878 | 176,505,885 | 88.86 |
| CM05843 | 237,168,860 | 210,884,888 | 88.92 |
| CM05844 | 218,006,446 | 193,777,063 | 88.89 |
| CM05847 | 193,898,986 | 171,620,068 | 88.51 |
| CM05849 | 195,102,658 | 173,127,054 | 88.74 |
| CM05853 | 207,074,456 | 183,763,775 | 88.74 |
| CM05854 | 203,654,150 | 180,196,467 | 88.48 |
| CM05855 | 207,188,646 | 183,903,644 | 88.76 |
| CM05856 | 227,767,066 | 203,309,224 | 89.26 |
| CM05857 | 209,752,072 | 185,743,392 | 88.55 |
| CM05858 | 228,902,536 | 203,126,606 | 88.74 |
| CM05863 | 211,808,824 | 186,837,922 | 88.21 |
| CM05865 | 201,372,348 | 178,523,693 | 88.65 |

| | | | |
|---|---|---|---|
| CM05869 | 233,540,912 | 206,812,444 | 88.56 |
| CM05870 | 234,807,068 | 207,595,324 | 88.41 |
| CM06496 | 216,046,576 | 192,510,081 | 89.11 |
| CM06501 | 215,915,406 | 191,622,423 | 88.75 |
| CM06505 | 220,509,940 | 194,982,858 | 88.42 |
| CM06510 | 197,621,370 | 174,879,247 | 88.49 |
| CM06511 | 211,953,450 | 187,963,418 | 88.68 |
| CM06513 | 211,394,978 | 187,598,559 | 88.74 |
| CM07226 | 228,566,352 | 202,935,403 | 88.79 |
| CM07234 | 226,123,688 | 200,571,269 | 88.70 |
| CM07244 | 226,518,412 | 198,905,704 | 87.81 |
| CM07246 | 215,005,996 | 189,602,042 | 88.18 |
| CM07249 | 275,946,300 | 238,679,461 | 86.49 |
| CM07258 | 235,161,112 | 207,433,380 | 88.21 |
| CM07261 | 210,081,902 | 185,331,317 | 88.22 |
| CM07262 | 261,031,932 | 230,130,021 | 88.16 |
| **CM07264** | 249,047,676 | 66,903,222 | 26.86 |
| CM07266 | 188,350,180 | 165,773,722 | 88.01 |
| CM07268 | 229,658,868 | 201,273,803 | 87.64 |
| **CM07269** | 190,691,498 | 47,256,832 | 24.78 |
| CM07270 | 257,206,834 | 226,383,379 | 88.02 |
| **CM07272** | 233,233,976 | 64,840,215 | 27.80 |
| **CM07277** | 223,332,526 | 55,224,057 | 24.73 |
| CM07280 | 185,520,622 | 163,378,721 | 88.06 |
| CM07281 | 250,905,090 | 220,355,900 | 87.82 |
| **CM07283** | 203,215,094 | 47,936,374 | 23.59 |
| **CM07285** | 181,737,890 | 49,206,639 | 27.08 |
| CM07289 | 182,189,834 | 160,286,764 | 87.98 |
| CM07292 | 174,182,552 | 152,118,994 | 87.33 |
| CM07295 | 201,397,410 | 177,211,916 | 87.99 |
| **CM07300** | 189,983,032 | 76,610,009 | 40.32 |
| CM07310 | 190,918,112 | 167,693,210 | 87.84 |
| CM07317 | 182,268,026 | 160,051,809 | 87.81 |
| CM07319 | 180,539,518 | 158,355,537 | 87.71 |
| CM07321 | 199,002,398 | 175,483,941 | 88.18 |
| CM07327 | 193,973,940 | 169,632,106 | 87.45 |
| CM07329 | 209,972,504 | 184,289,020 | 87.77 |
| CM07338 | 186,636,584 | 163,727,171 | 87.73 |
| CM07340 | 229,618,324 | 202,155,111 | 88.04 |
| CM07342 | 193,371,246 | 170,223,279 | 88.03 |
| CM07346 | 234,362,154 | 206,986,111 | 88.32 |

| | | | |
|---|---|---|---|
| CM07350 | 212,770,170 | 187,534,814 | 88.14 |
| CM07354 | 204,538,202 | 180,091,801 | 88.05 |
| CM07360 | 244,373,578 | 215,643,556 | 88.24 |
| CM07885 | 177,348,722 | 156,241,246 | 88.10 |
| CM07888 | 283,204,772 | 249,455,338 | 88.08 |
| CM07890 | 205,511,008 | 179,881,772 | 87.53 |
| CM07891 | 180,353,086 | 158,864,569 | 88.09 |
| CM07892 | 231,596,900 | 205,721,832 | 88.83 |
| CM07895 | 185,541,956 | 162,525,777 | 87.60 |
| CM07900 | 179,209,700 | 157,575,949 | 87.93 |
| CM07901 | 210,972,322 | 185,408,572 | 87.88 |
| CM07902 | 218,506,524 | 193,101,509 | 88.37 |
| CM07903 | 176,410,704 | 154,828,455 | 87.77 |
| CM07908 | 185,227,904 | 162,968,240 | 87.98 |
| CM07909 | 233,473,030 | 205,209,120 | 87.89 |
| CM08512 | 193,594,070 | 169,600,859 | 87.61 |
| CM08517 | 180,174,178 | 158,406,680 | 87.92 |
| CM08524 | 180,832,992 | 159,317,175 | 88.10 |
| CM08525 | 194,236,054 | 171,398,421 | 88.24 |
| CM08526 | 180,203,250 | 158,357,792 | 87.88 |
| CM08537 | 199,821,354 | 176,121,898 | 88.14 |
| CM08551 | 176,548,180 | 154,259,848 | 87.38 |
| CM08562 | 202,108,898 | 177,581,416 | 87.86 |
| CM08563 | 201,961,252 | 177,133,085 | 87.71 |
| CM08573 | 200,978,840 | 177,398,935 | 88.27 |
| CM08593 | 230,416,760 | 202,748,574 | 87.99 |
| CM08598 | 189,705,628 | 167,215,039 | 88.14 |
| CM08613 | 242,015,246 | 213,097,908 | 88.05 |
| CM08617 | 229,690,704 | 202,013,854 | 87.95 |
| CM08627 | 187,368,590 | 165,580,156 | 88.37 |
| CM08628 | 226,282,530 | 200,079,688 | 88.42 |
| CM08635 | 192,696,964 | 169,587,958 | 88.01 |
| CM08642 | 212,536,088 | 187,314,239 | 88.13 |
| CM08652 | 266,205,964 | 234,427,812 | 88.06 |
| CM08653 | 213,344,018 | 187,848,490 | 88.05 |
| CM08655 | 217,896,636 | 192,119,215 | 88.17 |
| CM08665 | 211,348,138 | 185,130,343 | 87.59 |
| CM08669 | 196,134,978 | 172,547,682 | 87.97 |
| CM08671 | 206,564,388 | 182,240,154 | 88.22 |
| CM08672 | 237,474,222 | 209,532,993 | 88.23 |
| CM08677 | 238,557,878 | 209,887,161 | 87.98 |

| | | | |
|---|---|---|---|
| CM08681 | 286,494,200 | 251,920,478 | 87.93 |
| CM08684 | 196,934,854 | 174,561,203 | 88.64 |
| CM12246 | 239,382,530 | 211,806,422 | 88.48 |
| CH1 | 225,012,214 | 198,839,767 | 88.37 |
| CH2 | 232,448,818 | 205,081,188 | 88.23 |
| CH3 | 248,177,072 | 219,250,259 | 88.34 |
| CH4 | 246,329,404 | 217,359,895 | 88.24 |
| 4349 | 75,582,392 | 36,744,544 | 48.62 |
| 67996 | 135,046,804 | 108,377,167 | 80.25 |
| If-Dlon1 | 119,875,036 | 95,345,810 | 79.54 |
| If-Dlon12 | 101,977,990 | 30,941,164 | 30.34 |
| L121 | 122,772,108 | 95,827,281 | 78.05 |
| L17 | 133,229,808 | 80,821,965 | 60.66 |
| L3027 | 114,951,968 | 88,884,130 | 77.32 |
| L4148 | 78,566,408 | 51,260,738 | 65.25 |
| L8249 | 72,275,374 | 60,238,085 | 83.35 |
| L9512 | 53,332,792 | 43,255,450 | 81.10 |
| L953 | 107,035,408 | 74,141,716 | 69.27 |
| M1070 | 97,130,012 | 69,725,636 | 71.79 |
| M861 | 108,392,832 | 30,872,258 | 28.48 |
| M865 | 90,924,046 | 54,193,455 | 59.60 |
| M883 | 116,617,722 | 88,168,368 | 75.60 |
| M905 | 103,620,152 | 86,784,457 | 83.75 |
| M949 | 135,802,752 | 113,724,806 | 83.74 |

**Supplementary Table 8.** Passport information of *D. exilis* and *D. longiflora* accessions. Included are accession name, country of origin, latitude (Lat.), longitude (Long.), altitude (Alt.), mean temperature of the wettest quarter (Mean temp.), mean precipitation of the wettest quarter (Mean prec.), ethnicity, linguistic groups and genetic cluster (based on $K$=6 and 70% ancestry threshold)

| Accession | Country | Lat. | Long. | Alt. | Mean temp. | Mean prec. | Ethnicity | Linguistic group | Genetic cluster |
|---|---|---|---|---|---|---|---|---|---|
| CM03380 | Togo | 10.30 | 0.42 | 121 | 25.8 | 667 | Chakosi | Volta-Congo | Admixed |
| CM03382 | Togo | 10.27 | 0.65 | 168 | 25.7 | 664 | Chakosi | Volta-Congo | 2 |
| CM03390 | Togo | 9.97 | 0.57 | 142 | 25.9 | 645 | Konkomba | Volta-Congo | 2 |
| CM03394 | Togo | 10.13 | 0.75 | 146 | 26 | 657 | Lamba | Volta-Congo | 2 |
| CM03396 | Togo | 10.03 | 0.92 | 213 | 25.5 | 677 | Lamba | Volta-Congo | 2 |
| CM03400 | Togo | 9.80 | 1.10 | 450 | 23.5 | 741 | Losso | Volta-Congo | 2 |
| CM03403 | Togo | 9.82 | 1.20 | 408 | 24.2 | 737 | Losso | Volta-Congo | 2 |
| CM03406 | Togo | 9.88 | 0.75 | 171 | 25.8 | 660 | Gangan | Volta-Congo | 2 |
| CM03409 | Togo | 10.02 | 0.83 | 223 | 25.5 | 671 | Lamba | Volta-Congo | 2 |
| CM03418 | Togo | 9.67 | 0.95 | 219 | 25.3 | 695 | Losso | Volta-Congo | 2 |
| CM03423 | Togo | 8.60 | 1.17 | 294 | 25 | 627 | Lamba | Volta-Congo | 5 |
| CM03424 | Togo | 9.57 | 0.70 | 202 | 25.5 | 662 | Konkomba | Volta-Congo | 5 |
| CM03430 | Togo | 9.27 | 0.80 | 281 | 25 | 685 | Bassari | Atlantic | 5 |
| CM03431 | Togo | 7.58 | 0.73 | 544 | 22.5 | 651 | Akposso | Volta-Congo | 5 |
| CM03434 | Togo | 7.62 | 1.02 | 418 | 23.5 | 607 | Akposso | Volta-Congo | 5 |
| CM03437 | Togo | 7.47 | 0.90 | 579 | 24.6 | 625 | Akposso | Volta-Congo | 5 |
| CM03438 | Togo | 9.42 | 0.92 | 328 | 24.7 | 693 | Kpelle | Mande Western | 5 |
| CM03439 | Togo | 6.57 | 0.85 | 118 | 27.2 | 466 | Kpelle | Mande Western | 5 |
| CM04487 | Benin | 10.50 | 1.38 | 383 | 25 | 692 | Ditamari | Volta-Congo | 2 |
| CM04489 | Benin | 10.40 | 0.78 | 184 | 25.8 | 662 | Niende | Volta-Congo | 2 |
| CM04493 | Benin | 10.27 | 0.98 | 203 | 26 | 657 | Ditamari | Volta-Congo | 2 |
| **CM04495** | Benin | 10.25 | 1.23 | 571 | 23.8 | 741 | NA | NA | Excluded |
| CM05734 | Mali | 13.62 | -7.03 | 329 | 26.2 | 517 | Bambara | Manding West | 6 |
| CM05736 | Mali | 13.88 | -7.18 | 308 | 26.5 | 491 | Sarakole | Mande Western | 6 |
| CM05737 | Mali | 14.02 | -7.52 | 358 | 26.3 | 494 | Bambara | Manding West | 6 |
| CM05740 | Mali | 14.87 | -10.30 | 256 | 27.5 | 494 | Bambara | Manding West | Admixed |
| CM05741 | Mali | 14.48 | -11.57 | 38 | 28.6 | 519 | Bambara | Manding West | 6 |
| CM05742 | Mali | 14.72 | -12.07 | 28 | 29 | 409 | Wolof | NA | 6 |
| CM05743 | Mali | 14.33 | -11.80 | 101 | 28.3 | 547 | Bambara | Manding West | 6 |
| CM05744 | Mali | 14.13 | -11.63 | 218 | 27.2 | 605 | Bambara | Manding West | 6 |
| CM05746 | Mali | 13.90 | -11.87 | 121 | 27.6 | 592 | Malinke | Manding West | 6 |
| CM05750 | Mali | 13.10 | -11.13 | 331 | 25.8 | 773 | Malinke | Manding West | Admixed |
| CM05754 | Mali | 12.27 | -10.88 | 261 | 25.7 | 888 | Malinke | Manding East | Admixed |
| **CM05757** | Mali | 12.82 | -10.33 | 263 | 26.2 | 790 | NA | NA | Excluded |
| CM05760 | Mali | 13.22 | -10.50 | 186 | 26.6 | 713 | Malinke | Manding West | Admixed |

| | | | | | | | | | |
|---|---|---|---|---|---|---|---|---|---|
| CM05761 | Mali | 13.20 | -10.75 | 139 | 26.9 | 722 | Malinke | Manding West | Admixed |
| CM05767 | Mali | 13.10 | -9.48 | 332 | 25.9 | 689 | Malinke | Manding West | Admixed |
| CM05770 | Mali | 12.97 | -9.22 | 333 | 26.1 | 712 | Bambara | Manding West | Admixed |
| CM05771 | Mali | 13.03 | -9.65 | 322 | 26 | 713 | Malinke | Manding West | Admixed |
| CM05772 | Mali | 12.88 | -9.95 | 331 | 26 | 754 | Malinke | Manding West | 4 |
| CM05775 | Mali | 13.88 | -8.10 | 398 | 26.3 | 571 | Bambara | Manding West | 6 |
| CM05780 | Mali | 13.10 | -7.93 | 391 | 25.9 | 680 | Bambara | Manding West | 6 |
| CM05792 | Mali | 10.60 | -8.20 | 411 | 25.4 | 827 | Fula | Atlantic | 4 |
| CM05793 | Mali | 11.03 | -8.22 | 378 | 25.6 | 777 | Fula | Atlantic | Admixed |
| CM05796 | Mali | 11.53 | -8.08 | 370 | 25.8 | 746 | Bambara | Manding West | 4 |
| CM05798 | Mali | 11.10 | -7.42 | 378 | 25.6 | 726 | Bambara | Manding West | Admixed |
| CM05802 | Mali | 10.47 | -7.45 | 379 | 25.5 | 799 | Bambara | Manding West | Admixed |
| CM05803 | Mali | 10.53 | -6.58 | 369 | 25.5 | 695 | Bambara | Manding West | Admixed |
| CM05806 | Mali | 10.82 | -6.88 | 354 | 25.7 | 690 | Bambara | Manding West | Admixed |
| CM05808 | Mali | 10.75 | -6.50 | 343 | 25.8 | 676 | Bambara | Manding West | 4 |
| CM05810 | Mali | 11.05 | -6.85 | 322 | 25.8 | 689 | Bambara | Manding West | 4 |
| CM05811 | Mali | 11.28 | -7.03 | 364 | 25.7 | 693 | Bambara | Manding West | 4 |
| CM05815 | Mali | 11.38 | -7.27 | 335 | 25.8 | 698 | Bambara | Manding West | Admixed |
| CM05819 | Mali | 10.75 | -6.13 | 338 | 25.8 | 712 | Senufo | Volta-Congo | 4 |
| CM05824 | Mali | 13.12 | -5.83 | 293 | 26.7 | 591 | Bambara | Manding West | Admixed |
| CM05827 | Mali | 12.65 | -4.90 | 314 | 26.7 | 596 | Minianka | Volta-Congo | Admixed |
| CM05830 | Mali | 13.60 | -4.60 | 275 | 27.2 | 493 | Bobo | NA | Admixed |
| CM05834 | Mali | 13.70 | -4.40 | 280 | 27.3 | 466 | Bobo | NA | Admixed |
| CM05835 | Mali | 14.02 | -4.23 | 270 | 27.5 | 404 | Marka | Mande Western | 1 |
| CM05836 | Mali | 15.05 | -3.88 | 263 | 27.9 | 333 | Fula | Atlantic | 1 |
| CM05836-D | Mali | 15.05 | -3.88 | 263 | 27.9 | 333 | Fula | Atlantic | Admixed |
| CM05839 | Mali | 14.28 | -2.55 | 269 | 28.3 | 379 | Dogon | Atlantic | 1 |
| CM05840 | Mali | 14.42 | -2.70 | 264 | 28.3 | 378 | Dogon | Atlantic | 1 |
| CM05841 | Mali | 14.57 | -2.90 | 293 | 28 | 379 | Dogon | Atlantic | 1 |
| CM05842 | Mali | 14.65 | -3.10 | 339 | 27.8 | 378 | Dogon | Atlantic | 1 |
| CM05843 | Mali | 14.13 | -3.62 | 388 | 27.3 | 405 | Dogon | Atlantic | 1 |
| CM05844 | Mali | 14.02 | -3.62 | 298 | 27.3 | 408 | Dogon | Atlantic | 1 |
| CM05847 | Mali | 13.75 | -3.95 | 388 | 26.4 | 438 | Dogon | Atlantic | 1 |
| CM05849 | Mali | 13.75 | -3.35 | 258 | 27.6 | 419 | Dogon | Atlantic | 1 |
| CM05853 | Mali | 14.30 | -3.37 | 351 | 27.1 | 399 | Dogon | Atlantic | 1 |
| CM05854 | Mali | 14.38 | -3.58 | 421 | 26.5 | 404 | Dogon | Atlantic | Admixed |
| CM05855 | Mali | 14.50 | -3.47 | 495 | 26.3 | 398 | Dogon | Atlantic | Admixed |
| CM05856 | Mali | 14.67 | -3.27 | 576 | 26.1 | 389 | Dogon | Atlantic | 1 |
| CM05857 | Mali | 14.33 | -3.77 | 403 | 26.9 | 400 | Dogon | Atlantic | 1 |
| CM05858 | Mali | 14.18 | -3.88 | 422 | 26.6 | 407 | Dogon | Atlantic | 1 |
| CM05863 | Mali | 13.58 | -6.00 | 288 | 26.8 | 508 | Sarakole | Mande Western | 6 |

| | | | | | | | | | |
|---|---|---|---|---|---|---|---|---|---|
| CM05865 | Mali | 13.65 | -5.55 | 278 | 26.6 | 542 | Bambara | Manding West | Admixed |
| CM05869 | Mali | 13.33 | -6.42 | 292 | 26.8 | 500 | Bambara | Manding West | 6 |
| CM05870 | Mali | 13.18 | -6.50 | 292 | 26.8 | 523 | Bambara | Manding West | 6 |
| CM06496 | Burkina Faso | 14.20 | -1.85 | 306 | 28.1 | 337 | Fula | NA | 1 |
| CM06501 | Burkina Faso | 13.37 | -3.08 | 272 | 27.4 | 437 | Samo | Mande Eastern | 1 |
| CM06505 | Burkina Faso | 12.82 | -3.62 | 259 | 27.1 | 488 | Dafi | Volta-Congo | Admixed |
| CM06510 | Burkina Faso | 12.90 | -3.90 | 274 | 27 | 512 | Bobo | Mande Western | Admixed |
| CM06511 | Burkina Faso | 12.67 | -3.83 | 267 | 26.9 | 527 | Bobo | Mande Western | Admixed |
| CM06513 | Burkina Faso | 12.58 | -3.82 | 268 | 27 | 538 | Bobo | Mande Western | Admixed |
| CM07226 | Guinea | 11.07 | -9.25 | 355 | 25.5 | 874 | Malinke | Manding East | Admixed |
| CM07234 | Guinea | 10.87 | -9.50 | 392 | 25.2 | 907 | Malinke | Manding East | 4 |
| CM07244 | Guinea | 10.63 | -10.97 | 496 | 25.1 | 851 | Fula | Atlantic | 3 |
| CM07246 | Guinea | 10.72 | -11.22 | 605 | 24.7 | 868 | Malinke | Manding East | 4 |
| CM07249 | Guinea | 10.70 | -11.50 | 868 | 23.1 | 954 | Fula | Atlantic | 3 |
| CM07258 | Guinea | 10.90 | -11.12 | 424 | 25.5 | 835 | Fula | Atlantic | Admixed |
| CM07261 | Guinea | 11.22 | -12.13 | 903 | 23.4 | 1035 | Fula | Atlantic | Admixed |
| CM07262 | Guinea | 10.64 | -12.33 | 955 | 20.2 | 1211 | Fula | Atlantic | Admixed |
| **CM07264** | Guinea | 10.73 | -12.12 | 791 | 22.5 | 1082 | NA | NA | Excluded |
| CM07266 | Guinea | 10.83 | -12.50 | 638 | 21.8 | 1183 | Fula | Atlantic | Admixed |
| CM07268 | Guinea | 11.38 | -12.13 | 858 | 22.8 | 1028 | Fula | Atlantic | 4 |
| **CM07269** | Guinea | 11.30 | -11.87 | 768 | 22.8 | 1031 | NA | NA | Excluded |
| CM07270 | Guinea | 11.30 | -11.87 | 751 | 22.8 | 1031 | Fula | Atlantic | Admixed |
| **CM07272** | Guinea | 11.53 | -11.57 | 727 | 23.6 | 997 | NA | NA | Excluded |
| **CM07277** | Guinea | 11.50 | -11.87 | 792 | 22.9 | 1028 | NA | NA | Excluded |
| CM07280 | Guinea | 11.72 | -11.98 | 622 | 24.1 | 993 | Fula | Atlantic | Admixed |
| CM07281 | Guinea | 11.57 | -12.15 | 738 | 23.4 | 1024 | Fula | Atlantic | Admixed |
| **CM07283** | Guinea | 11.57 | -12.15 | 750 | 23.4 | 1024 | NA | NA | Excluded |
| **CM07285** | Guinea | 11.45 | -12.23 | 884 | 22.5 | 1034 | NA | NA | Excluded |
| CM07289 | Guinea | 11.87 | -12.37 | 854 | 22.8 | 1043 | Fula | Atlantic | Admixed |
| CM07292 | Guinea | 12.03 | -12.50 | 686 | 23.2 | 1033 | Fula | Atlantic | Admixed |
| CM07295 | Guinea | 11.58 | -12.43 | 994 | 21.7 | 1080 | Fula | Atlantic | Admixed |
| **CM07300** | Guinea | 12.17 | -13.07 | 149 | 26.2 | 1040 | NA | NA | Excluded |
| CM07310 | Guinea | 11.30 | -13.18 | 181 | 25.4 | 1356 | Fula | Atlantic | Admixed |
| CM07317 | Guinea | 10.60 | -13.15 | 346 | 25 | 1549 | Fula | Atlantic | Admixed |
| CM07319 | Guinea | 10.33 | -12.95 | 235 | 25.5 | 1488 | Susu | Mande Western | Admixed |
| CM07321 | Guinea | 10.00 | -12.68 | 166 | 25.6 | 1397 | Susu | Mande Western | Admixed |
| CM07327 | Guinea | 10.23 | -11.97 | 719 | 22.2 | 1086 | Fula | Atlantic | 3 |
| CM07329 | Guinea | 10.13 | -11.63 | 393 | 24.7 | 982 | Fula | Atlantic | 3 |
| CM07338 | Guinea | 9.53 | -10.52 | 529 | 24.3 | 918 | Malinke | Manding East | 4 |
| CM07340 | Guinea | 9.40 | -10.22 | 520 | 24.4 | 936 | Malinke | Manding East | 4 |
| CM07342 | Guinea | 9.35 | -9.97 | 493 | 24.6 | 964 | Malinke | Manding East | 4 |

| | | | | | | | | | |
|---|---|---|---|---|---|---|---|---|---|
| CM07346 | Guinea | 9.72 | -9.77 | 465 | 24.7 | 942 | Malinke | Manding East | 4 |
| CM07350 | Guinea | 10.00 | -9.15 | 417 | 25.2 | 933 | Malinke | Manding East | 4 |
| CM07354 | Guinea | 9.03 | -9.00 | 537 | 24.1 | 998 | Malinke | Manding East | 4 |
| CM07360 | Guinea | 10.28 | -8.33 | 446 | 25.1 | 878 | Malinke | Manding East | 4 |
| CM07885 | Burkina Faso | 10.55 | -5.28 | 360 | 25.3 | 694 | Senufo | Volta-Congo | 2 |
| CM07888 | Burkina Faso | 10.55 | -5.08 | 313 | 25.8 | 682 | Gouin | Volta-Congo | Admixed |
| CM07890 | Burkina Faso | 10.58 | -4.85 | 290 | 26.1 | 682 | Dyula | Volta-Congo | 2 |
| CM07891 | Burkina Faso | 10.63 | -4.90 | 297 | 26 | 678 | Turka | Volta-Congo | Admixed |
| CM07892 | Burkina Faso | 10.67 | -4.82 | 285 | 26.3 | 674 | Karaboro | Volta-Congo | 2 |
| CM07895 | Burkina Faso | 10.52 | -4.98 | 303 | 25.9 | 685 | Turka | Volta-Congo | Admixed |
| CM07900 | Burkina Faso | 11.18 | -4.92 | 426 | 25.5 | 674 | Toussian | Volta-Congo | 2 |
| CM07901 | Burkina Faso | 11.05 | -4.93 | 534 | 24.7 | 681 | Siamou | Volta-Congo | Admixed |
| CM07902 | Burkina Faso | 11.18 | -4.62 | 424 | 25.4 | 660 | Sambla | Mande Western | Admixed |
| CM07903 | Burkina Faso | 11.13 | -4.43 | 388 | 25.6 | 658 | Bobo | Mande Western | 2 |
| CM07908 | Burkina Faso | 11.13 | -4.33 | 460 | 25.2 | 667 | Toussian | Mande Western | 2 |
| CM07909 | Burkina Faso | 11.02 | -4.05 | 316 | 26 | 638 | Tiefo | Volta-Congo | Admixed |
| CM08512 | Guinea | 10.24 | -12.03 | 672 | 22.6 | 1090 | Fula | Atlantic | Admixed |
| CM08517 | Guinea | 10.69 | -12.25 | 1122 | 19.9 | 1205 | Fula | Atlantic | Admixed |
| CM08524 | Guinea | 11.20 | -12.07 | 701 | 23.3 | 1030 | Sarakole | Mande Western | Admixed |
| CM08525 | Guinea | 10.80 | -12.21 | 1036 | 20.5 | 1161 | Fula | Atlantic | Admixed |
| CM08526 | Guinea | 11.01 | -12.05 | 695 | 23.5 | 1038 | Fula | Atlantic | Admixed |
| CM08537 | Guinea | 10.88 | -12.18 | 697 | 23.2 | 1073 | Fula | Atlantic | 4 |
| CM08551 | Guinea | 11.32 | -12.29 | 1036 | 21.5 | 1030 | Fula | Atlantic | Admixed |
| CM08562 | Guinea | 11.40 | -12.40 | 1008 | 21.5 | 1064 | Fula | Atlantic | Admixed |
| CM08563 | Guinea | 11.34 | -12.50 | 898 | 21.6 | 1104 | Fula | Atlantic | Admixed |
| CM08573 | Guinea | 11.65 | -12.35 | 525 | 23.9 | 1045 | Fula | Atlantic | Admixed |
| CM08593 | Guinea | 11.35 | -11.93 | 759 | 23 | 1025 | Fula | Atlantic | Admixed |
| CM08598 | Guinea | 11.28 | -10.71 | 424 | 25.3 | 886 | Fula | Atlantic | Admixed |
| CM08613 | Guinea | 10.72 | -10.71 | 580 | 25.2 | 868 | Malinke | Manding East | 4 |
| CM08617 | Guinea | 10.85 | -10.94 | 420 | 25.6 | 856 | Fula | Atlantic | Admixed |
| CM08627 | Guinea | 10.79 | -10.18 | 417 | 25.1 | 895 | Malinke | Manding East | 4 |
| CM08628 | Guinea | 10.71 | -9.58 | 367 | 25.2 | 917 | Malinke | Manding East | 4 |
| CM08635 | Guinea | 9.88 | -9.55 | 434 | 25.2 | 939 | Malinke | Manding East | 4 |
| CM08642 | Guinea | 9.63 | -9.07 | 510 | 24.7 | 956 | Malinke | Manding East | 4 |
| CM08652 | Guinea | 10.04 | -10.75 | 443 | 24.9 | 827 | Malinke | Manding East | 4 |
| CM08653 | Guinea | 9.96 | -10.70 | 458 | 25 | 834 | Malinke | Manding East | 4 |
| CM08655 | Guinea | 9.20 | -10.58 | 661 | 23.6 | 996 | Malinke | Manding East | 4 |
| CM08665 | Guinea | 9.83 | -13.11 | 342 | 25 | 1931 | Susu | Mande Western | 3 |
| CM08669 | Guinea | 9.99 | -12.91 | 391 | 24.1 | 1460 | Susu | Mande Western | 4 |
| CM08671 | Guinea | 9.83 | -13.21 | 222 | 26 | 2173 | Susu | Mande Western | Admixed |
| CM08672 | Guinea | 9.97 | -12.55 | 146 | 25.9 | 1342 | Fula | Atlantic | Admixed |

| | | | | | | | | | |
|---|---|---|---|---|---|---|---|---|---|
| CM08677 | Guinea | 9.79 | -13.35 | 241 | 25.4 | 2379 | Susu | Mande Western | Admixed |
| CM08681 | Guinea | 9.58 | -13.29 | 20 | 26.5 | 2467 | Susu | Mande Western | Admixed |
| CM08684 | Guinea | 10.03 | -13.65 | 37 | 27.1 | 2457 | Susu | Mande Western | Admixed |
| CM12246 | Niger | 13.04 | 3.22 | 239 | 28 | 430 | NA | NA | Admixed |
| CH1 | Ghana | 10.16 | 0.26 | 190 | 25.5 | 665 | NA | NA | 2 |
| CH2 | Ghana | 10.16 | 0.26 | 190 | 25.5 | 665 | NA | NA | 2 |
| CH3 | Ghana | 10.13 | 0.29 | 174 | 25.5 | 665 | NA | NA | 2 |
| CH4 | Ghana | 10.14 | 0.29 | 160 | 25.6 | 665 | NA | NA | 2 |

***Digitaria longiflora***

| **Accession** | **Country** | **Latitude** | **Longitude** | **Altitude** | **Mean temp.** | **Mean prec.** | **Ethnicity** | **Linguistic group** | **Group** |
|---|---|---|---|---|---|---|---|---|---|
| 4349 | NA | NA | NA | NA | NA | NA | NA | NA | wild |
| 67996 | Guinea | 7.70 | -8.40 | 734 | 21.7 | 933 | NA | NA | wild |
| If-Dlon1 | Mali | 14.58 | -4.23 | 268 | 27.7 | 391 | NA | NA | wild |
| If-Dlon12 | Sierra Leone | 9.18 | -11.12 | 1104 | 20.3 | 1155 | NA | NA | wild |
| L121 | South Sudan | 6.20 | 31.68 | 433 | 25.7 | 402 | NA | NA | wild |
| L17 | Nigeria | 6.45 | 3.43 | 11 | 26.5 | 899 | NA | NA | wild |
| L3027 | Chad | 11.63 | 16.03 | 327 | 27.6 | 444 | NA | NA | wild |
| L4148 | Nigeria | 9.75 | 10.50 | 284 | 25.7 | 640 | NA | NA | wild |
| L8249 | Guinea | 7.70 | -8.40 | 745 | 21.7 | 933 | NA | NA | wild |
| L9512 | Cameroon | 4.97 | 9.93 | 921 | 20.4 | 1267 | NA | NA | wild |
| L953 | Kenya | -0.62 | 34.67 | 1516 | 20.7 | 632 | NA | NA | wild |
| M1070 | Nigeria | 7.38 | 3.94 | 220 | 25.7 | 523 | NA | NA | wild |
| M861 | NA | NA | NA | NA | NA | NA | NA | NA | wild |
| M865 | NA | NA | NA | NA | NA | NA | NA | NA | wild |
| M883 | Cameroon | 5.73 | 10.90 | 1137 | 21.2 | 915 | NA | NA | wild |
| M905 | Chad | 14.70 | 18.59 | 330 | 30.2 | 153 | NA | NA | wild |
| M949 | Chad | 14.70 | 18.59 | 330 | 30.2 | 153 | NA | NA | wild |

**Supplementary Table 9**. Analysis of covariance (ANCOVA) for testing the effect of all factors together.

| Variables | Df | Sum Sq | Mean Sq | F value | Pr(>F) |
|---|---|---|---|---|---|
| Temperature | 1 | 4273340 | 4273340 | 306.145 | < 2.2e-16 |
| Precipitation | 1 | 1138326 | 1138326 | 81.5505 | 7.41E-15 |
| Latitude | 1 | 297866 | 297866 | 21.3393 | 1.07E-05 |
| Longitude | 1 | 1084859 | 1084859 | 77.7201 | 2.26E-14 |
| Altitude | 1 | 1260 | 1260 | 0.0903 | 0.76442 |
| Ethnic | 31 | 683156 | 22037 | 1.5788 | 0.04503 |
| Linguistic | 3 | 64209 | 21403 | 1.5333 | 0.21003 |

**Supplementary Table 10.** Origin of wild *D. longiflora* accessions used in this study.

| Accession | Country | Collection |
|---|---|---|
| 4349 | NIGERIA | Cirad |
| 67996 | GUINEA | Cirad |
| If-Dlon1 | MALI | IFAN Ch. A. Diop |
| If-Dlon12 | SIERRA LEONE | IFAN Ch. A. Diop |
| L121 | SOUDAN | National Herbarium of The Netherlands |
| L17 | NIGERIA | National Herbarium of The Netherlands |
| L3027 | TCHAD | National Herbarium of The Netherlands |
| L4148 | NIGERIA | National Herbarium of The Netherlands |
| L8249 | GUINEA | National Herbarium of The Netherlands |
| L9512 | CAMEROUN | National Herbarium of The Netherlands |
| L953 | KENYA | National Herbarium of The Netherlands |
| M1070 | NIGERIA | Museum of Natural History of Paris |
| M861 | GABON | Museum of Natural History of Paris |
| M865 | CONGO | Museum of Natural History of Paris |
| M883 | CAMEROUN | Museum of Natural History of Paris |
| M905 | TCHAD | Museum of Natural History of Paris |
| M949 | TCHAD | Museum of Natural History of Paris |

**Supplementary Table 11.** Alignment of 18 largest Hi-C super-scaffolds (DeNovoMAGIC3 + Hi-C) to the respective hybrid scaffolds (DeNovoMAGIC3 + optical map). The hybrid scaffolds highlighted in red detected chimeras in the original DeNovoMAGIC3 assembly.

| Hi-C super- scaffold | Hybrid scaffold |
|---|---|
| 1059 | 100113 / 444 |
| 8292 | 29 / 106 |
| 233 | 274 / 100007 |
| 8290 | 418 / 528 |
| 208 | 375 / 262 |
| 94 | 100001 / 100018 |
| 2 | 100020 / 100042 / 100043 |
| 103 | 100024 / 100028 |
| 8291 | 100015 / 79 |
| 1050 | 100010 / 528 |
| 190 | 100030 / 103 |
| 52 | 100016 / 248 |
| 8293 | 100004 / 103 |
| 86 | 100045 / 219 |
| 8288 | 138 / 100019 |
| 139 | 8894 / 100051 / 100033 |
| 8289 | 100005 / 100002 |
| 117 | 100003 / 16986 |

**Supplementary Table 12.** Number of SNPs retained after each filtering step.

| Filtering step | Number of SNPs retained |
|---|---|
| 1- Initial VCF | 36,514,446 |
| 2- SNP clusters | 24,721,046 |
| 3- InDel removal | 17,471,641 |
| 4- Variant missingness | 15,556,597 |
| 5- Mean depth | 15,378,274 |
| 6- Biallelic SNPs | 14,076,228 |
| 7- Missing data per accession | 11,517,157 |
| 8- SNPs in unanchored chromosome | 11,046,501 |
